# Obstacles to Studying Alternative Splicing Using scRNA-seq

**DOI:** 10.1101/797951

**Authors:** Jennifer Westoby, Pavel Artemov, Martin Hemberg, Anne Ferguson-Smith

## Abstract

**Background:** Early single-cell RNA-seq (scRNA-seq) studies suggested that it was unusual to see more than one isoform being produced from a gene in a single cell, even when multiple isoforms were detected in matched bulk RNA-seq samples. However, these studies generally did not consider the impact of dropouts or isoform quantification errors, potentially confounding the results of these analyses.

**Results:** In this study, we take a simulation based approach in which we explicitly account for dropouts and isoform quantification errors. We use our simulations to ask to what extent it is possible to study alternative splicing using scRNA-seq. Additionally, we ask what limitations must be overcome to make splicing analysis feasible. We find that the high rate of dropouts associated with scRNA-seq is a major obstacle to studying alternative splicing. In mice and other well established model organisms, the relatively low rate of isoform quantification errors poses a lesser obstacle to splicing analysis. We find that different models of isoform choice meaningfully change our simulation results.

**Conclusions:** To accurately study alternative splicing with single-cell RNA-seq, a better understanding of isoform choice and the errors associated with scRNA-seq is required. An increase in the capture efficiency of scRNA-seq would also be beneficial. Until some or all of the above are achieved, we do not recommend attempting to resolve isoforms in individual cells using scRNA-seq.

## Background

Single-cell RNA-seq (scRNA-seq) theoretically enables transcriptomic analysis at single cell resolution. If measurements are accurate, the data would allow fundamental molecular biology questions regarding how alternative splicing is regulated at the cellular level to be addressed. However, to date the majority of scRNA-seq studies have been analysed at the gene rather than the transcript level. Isoform quantification remains a challenging problem for bulk RNA-seq [1, 2] and we suspect that many researchers are concerned that the high degree of technical noise associated with scRNA-seq could overwhelm any biological signal from alternative splicing events. Effectively distinguishing between technical and biological noise is made all the more challenging by a lack of orthogonal methods for validating scRNA-seq. Single molecule FISH (smFISH) can be used to validate some of the predictions of scRNA-seq, but resolving between highly similar isoforms remains challenging [3, 4, 5, 6]. Although the throughput of smFISH is improving [7], to the best of our knowledge no smFISH technology currently exists which could accurately resolve a high proportion of the transcriptome at an isoform level.

A recent benchmark of isoform quantification for scRNA-seq found that many isoform quantification softwares perform almost as well for scRNA-seq datasets as for bulk RNA-seq [8]. Whilst this is encouraging, it is important to note that the benchmark only evaluated the ability of quantification softwares to correctly assign simulated scRNA-seq reads to the transcripts that generated them. It is well known that a substantial amount of scRNA-seq technical noise occurs prior to the bioinformatic analysis of reads, most notably dropouts due to a low capture efficiency and PCR amplification bias due to a low amount of starting material [9, 10, 11]. To the best of our knowledge, the impact of these and other sources of technical noise on splicing analysis accuracy in scRNA-seq experiments has not been systematically studied. Consequently, the extent to which it is possible to accurately perform splicing analysis with scRNA-seq is not well understood.

It has been known for some time that not all isoforms are equally likely to be expressed from a given gene. Bulk RNA-seq studies have shown that most genes have a ‘major’, highly abundant isoform, and will sometimes additionally have ‘minor’, more lowly expressed isoforms [12, 13]. It is not currently understood how isoform choice is regulated at the cellular level for most genes. In particular, it is not clear whether all cells express all isoforms but at different levels, or whether each cell exclusively express one or a subset of the total number of possible isoforms for a given gene. Such knowledge has the potential to contribute to a greater understanding of splicing mechanisms. For example, it is not known to what extent a common mechanism might be used to regulate isoform number at a cellular level, or whether every gene is substantially different. Several scRNA-seq studies have found that for genes which expressed multiple isoforms in bulk RNA-seq, only one or a small number of isoforms were detected in matched scRNA-seq [14, 15, 16, 17]. However, many of these studies did not consider the impact of dropouts and quantification software errors, potentially confounding their conclusions. The deceptively simple question: ‘How many isoforms are produced from a gene in a single cell?’ has a central place in our understanding of molecular biology, yet its answer remains unclear.

In this study, we return to this basic biological question using a fundamentally different approach. We take real scRNA-seq datasets and select genes for which four isofoms are detected. We then use these genes to simulate the following four scenarios: 1) all cells express one isoform per gene per cell, 2) all cells express two isoforms per gene per cell, 3) all cells express three isoforms per gene per cell, and 4) all cells express four isoforms per gene per cell. Importantly, in each scenario we explicitly simulate dropout events and quantification errors. We then use the simulated output of each scenario to ask two questions. Firstly, to what extent are we able to distinguish between these global differences in alternative splicing using scRNA-seq? And secondly, what should be done to enable more accurate splicing analysis with scRNA-seq?

## Results

A detailed description of our simulation approach can be found in Methods, a brief description is given here for convenience. Our approach for the first scenario, in which we simulate one isoform being expressed per gene per cell, is to first identify genes for which the expression of exactly four isoforms is detected in a real scRNA-seq dataset. In the second step, we randomly select one isoform based on a plausible model of isoform choice for the first of our genes in the first cell in our simulated dataset. For our default model of isoform choice, we choose the isoform based on a model of alternative splicing described by Hu et al [18]. Third, we simulate dropouts based on a Michaelis-Menten model described by Andrews and Hemberg [9]. Fourth, we simulate quantification errors based on isoform detection error estimates based on work by Westoby et al [8]. We repeat these four steps for every four isoform gene and cell in our simulated dataset, then calculate the mean number of isoforms detected for that gene per cell. The entire process described above is one complete simulation. We run 100 simulations for each of our four scenarios, where each scenario corresponds to one, two, three or four isoforms being expressed per gene per cell. We can then plot the distributions of the mean number of isoforms detected per gene per cell for each scenario.

In Figure 1, we apply our simulation approach to a dataset of H1 and H9 human embryonic stem cells (hESCs) [19]. In this dataset, each cell’s cDNA was split into two groups and sequenced at two different sequencing depths, enabling us to directly compare our simulation results at different sequencing depths without biological confounders. One group was sequenced at approximately 1 million reads per cell and the other group at approximately 4 million reads per cell on average. Our simulation results for the two H1 groups are compared side by side in Figure 1a. scRNA-seq experiments have been found to saturate in terms of the number of genes detected per cell at approximately 1 million reads per cell [20, 21]. However, we observe differences in the number of isoforms detected per gene per cell at 1 and 4 million reads per cell, indicating that the saturation depth may differ for gene and isoform level analyses.

**Figure 1:**
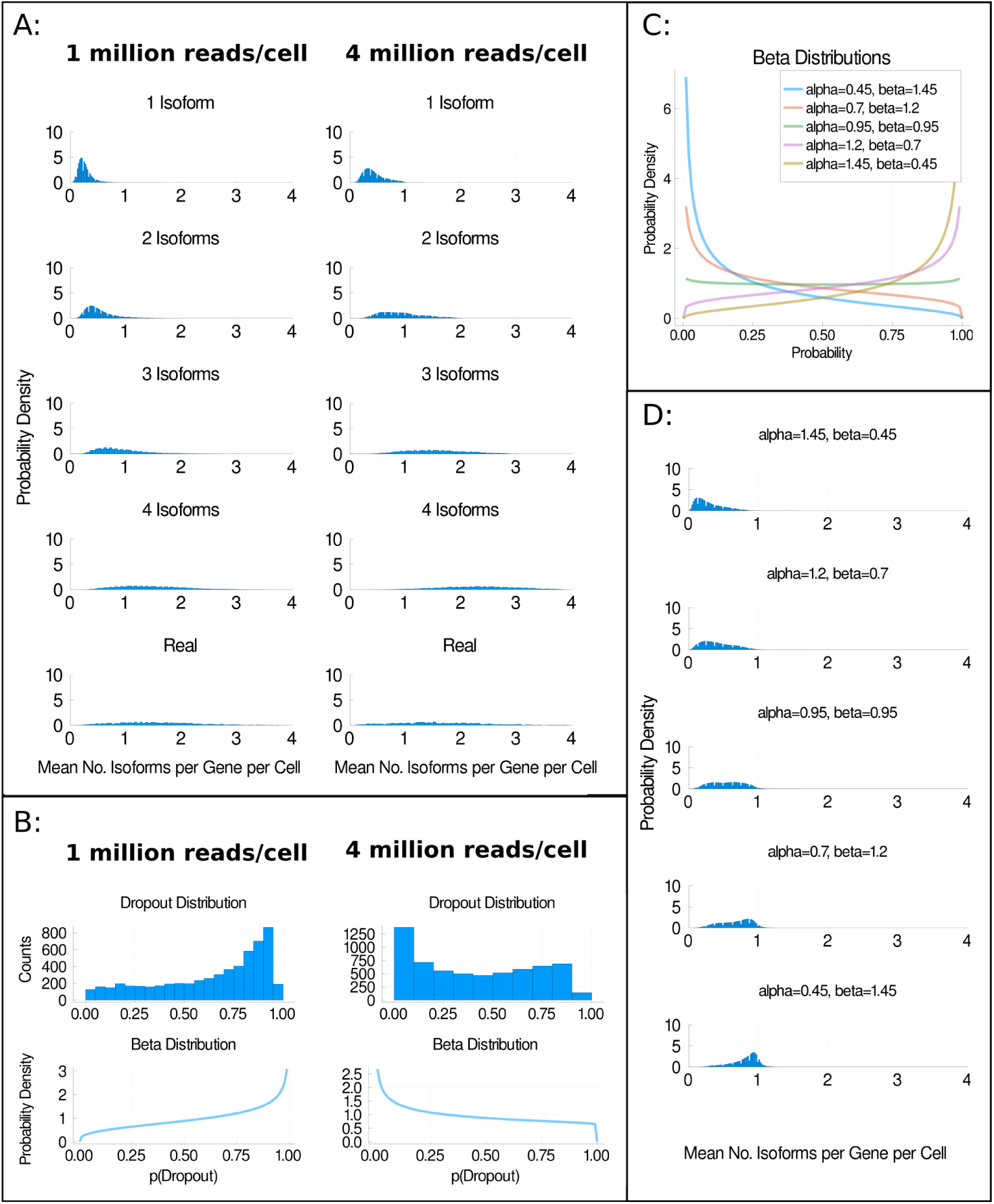
The impact of dropouts on isoform detection. **a** Distributions of the mean number of isoforms detected per gene per cell for H1 hESCs whose cDNA was split and sequenced at approximately 1 million reads per cell or 4 million reads per cell on average. **b** shows the distribution of the probabilities of dropouts (p(Dropout)) in each group and an approximation of these distributions using a Beta distribution. At 1 million reads per cell, *α* = 1.31 and *β* = 0.74 in the approximated Beta distribution. At 4 million reads per cell, *α* = 0.72 and *β* = 1.03 in the approximated Beta distribution. **c** shows five Beta Distributions from which dropout probabilities were sampled from inf the simulations used to generate **d**. In **d**, the distribution of the mean number of isoforms detected per gene per cell is shown for simulations in which one isoform was produced per gene per cell. Each plot corresponds to a simulation in which dropout probabilities were sampled from one of the distributions shown in **c**. Plots shown in **d** are for H1 hESCs sequenced at 4 million reads per cell. See Supplementary Figure 1 for equivalent plots for the H9 hESCs.

Figure 1a illustrates some of the difficulties associated with splicing analysis in scRNA-seq. At both sequencing depths, the distributions of the observed mean number of isoforms per gene per cell are shifted substantially to the left of their true value. This effect is less extreme, but still present, for the group sequenced at approximately 4 million reads per cell compared to the group sequenced at 1 million reads per cell. This is consistent with the hypothesis that sequencing at higher depth reduces the extent to which isoform number is underestimated. However, even at approximately 4 million reads per cell our simulations suggest that scRNA-seq substantially underestimates the mean number of isoforms per gene per cell for almost all genes. A naive analysis of these two datasets would most likely underestimate the number of isoforms expressed per gene per cell. This casts doubt on the biological relevance of previous observations suggesting only one isoform was typically produced per gene per cell, although admittedly the sequencing depth per cell was generally much greater than 4 million reads per cell in those studies (for example, Shalek et al. sequenced approximately 27 million reads per cell [14]).

One hypothesis for why our ability to detect isoforms increases with increased sequencing depth is that the rate of dropouts is reduced. In Figure 1b, we investigate this hypothesis by plotting the distribution of the probabilities of dropout for each isoform (p(dropout)), as estimated using the Michaelis-Menten equation [9] (see Methods). We find that the distribution is skewed towards high probabilities of dropout for the group sequenced at around 1 million reads per cell. In contrast, the distribution for the group sequenced at around 4 million reads per cell is more skewed towards low probabilities of dropouts. This demonstrates that our estimated dropout probabilities are different at the two sequencing depths, as expected.

Overall, the data in Figure 1a and 1b support the hypothesis that when the rate of technical dropouts decreases, the accuracy of isoform number estimation increases. However, as our dataset was only sequenced at two depths, we only have two data points available to investigate our hypothesis. To extend our investigation, we assume that the distributions of dropout probabilities observed in Figure 1b can be modelled as Beta distributions. The Beta distribution is parameterised by two values, *α* and *β*, and we find that it approximates our probability distributions well (see bottom panels of Figure 1b). Therefore, we select five values of *α* and *β* that generate differently shaped dropout distributions, as shown in Figure 1c. We then perform five further simulation experiments. In each simulation experiment, we sample our dropout probabilities from one of our Beta distributions. The results of these experiments are shown in Figure 1d.

In Figure 1d, we show the mean detected number of isoforms per gene per cell for the scenario where each gene produces one isoform per gene per cell. As we move from the top to the bottom of Figure 1d, the value of *α* decreases, corresponding to scenarios where the probability of dropout is more frequently close to zero. As *α* decreases, the distributions of mean detected isoforms per gene per cell shift further to the right and closer to the true number of isoforms produced per cell. We conclude that reducing the dropout rate would likely improve the accuracy of splicing analyses performed using scRNA-seq.

### Quantification errors are a relatively minor obstacle to studying alternative splicing

A benchmark of isoform quantification softwares in full length coverage mouse scRNA-seq datasets found that the error rate of many software tools was low and comparable to bulk RNA-seq [8]. This is encouraging, however it should be noted that the error rate is likely to be substantially higher for non-model organisms with less well annotated genomes than the mouse genome. As isoform quantification is a key step of many scRNA-seq alternative splicing analysis pipelines, it would be beneficial understand how quantification errors impact our ability to study alternative splicing, both when the error rate is high and when the error rate is low.

As our interest in this study is the detected number of isoforms per gene per cell, we are only interested in quantification errors which lead to changes in the number of isoforms detected. We simulate two types of quantification errors, false positives and false negatives. In this context, a false positive occurs when an isoform is called as expressed by the quantification software when there are no reads from that isoform. Note that this means that if an isoform is expressed in a cell but no reads are captured from it (i.e. a dropout), but the quantification software calls it as expressed, we would define this as a false postive event. A false negative occurs when an isoform is not called as expressed by the isoform quantification software when reads from that isoform are present. Based on our previous benchmark [8], we estimate that the probability of false positive events (*pFP*) is around 1% and that the probability of false negative (*pFN*) events is around 4% (see Methods). In our simulations in Figure 2, we vary both of these probabilities in the range of 0% to 50%. Figure 2a shows how the mean number of isoforms detected per gene per cell distributions changes as the probability of false positives and false negatives alters when every gene expresses one isoform per cell. Importantly, even when the probability of false positives and false negatives is zero, the peak of the mean number of detected isoforms per gene per cell is substantially shifted from its true value of one. This indicates that even if a perfect, 100% accurate isoform quantification tool existed, there would still be substantial barriers to studying alternative splicing using scRNA-seq. We suspect that the reason a 100% accurate isoform quantification tool would underestimate the number of isoforms per gene per cell is that isoform quantification tools usually only quantify the reads that are present. Due to the high number of dropouts in scRNA-seq, many expressed isoforms do not generate reads and thus would be called as unexpressed by a 100% accurate isoform quantification tool, leading to an under-estimate of the number of isoforms present.

**Figure 2:**
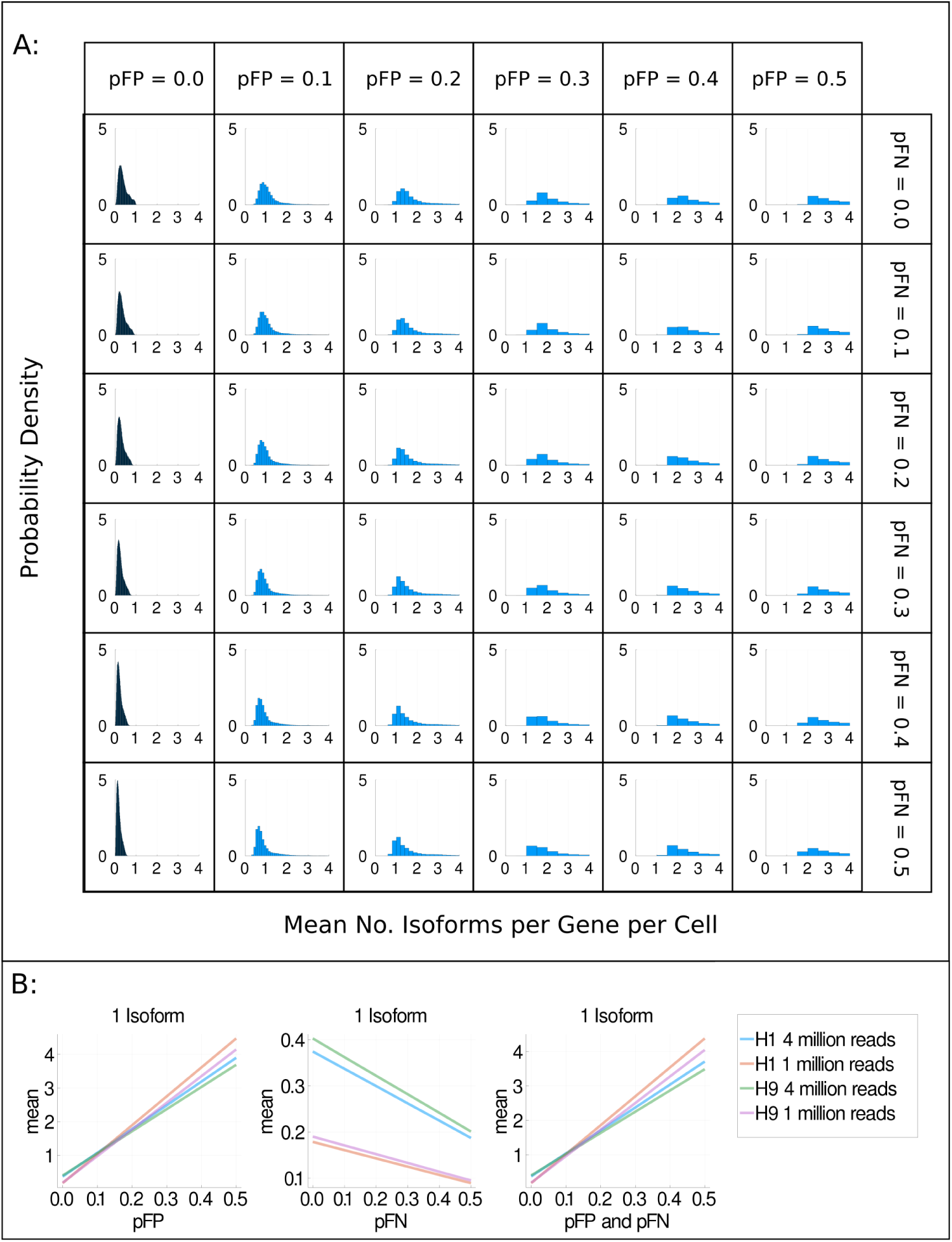
The impact of quantification errors on isoform detection. **a** Distributions of the mean number of isoforms detected per gene per cell when one isoform is expressed per gene per cell. The probability of false positives (*pFP*) increases from left to right and the probability of false negatives (*pFN*) increases from top to bottom. The dataset shown is H1 hESCs whose cDNA was split and sequenced at approximately 4 million reads per cell on average. **b** Summary plots of the average of the mean number of isoforms detected per gene per cell when *pFP, pFN*, or *pFP* and *pFN* are increased.

Unsurprisingly, increasing the probability of false positives causes an increase in the mean number of detected isoforms whilst increasing the probability of false negatives causes the mean number of detected isoforms to decrease, as shown in Figure 2b. Somewhat counterintuitively, increasing the probability of false positives from 0.0 to 0.1 actually shifts the peak of the mean isoforms detected distribution closer to its true value of one. This is probably because slightly increasing the probability of false positives allows some dropout events to be detected. Even though the number of isoforms detected is correct, the identities of many of the isoforms called as expressed are probably incorrect. Similarly, a high proportion of expressed isoforms are likely not detected due to dropouts. Interestingly, when the probability of false positives and false negatives are equally increased (the diagonal of Figure 2a), the mean number of detected isoforms increases, suggesting that the increased rate of false positives dominates over the increased rate of false negatives. This is likely because more isoforms are unexpressed than are expressed, and thus there are more opportunities for false positive events than for false negative events. Overall, we find that high probabilities of false positives and false negatives decrease our ability to accurately detect expressed isoforms in scRNA-seq.

### Different models of isoform choice meaningfully change our simulation results

It is possible that different mechanisms of isoform choice at the cellular level could alter our ability to correctly detect what isoforms are present in scRNA-seq. Because there is uncertainty over the mechanism of isoform choice within single cells, we implement four different models of isoform choice in our simulations. We then ask whether different models of isoform choice alter the mean number of detected isoforms per gene per cell in our simulations.

We give a detailed description of how each of these models was implemented in the Methods section, here we provide a brief description of each model and the rationale behind it. We first model the alternative splicing process as a type III Weibull distribution, using a model described by Hu et al [18]. Based on observations about the molecular process of alternative splicing, Hu et al suggested that the process could be well modelled by an extreme value distribution, and they found that a Weibull distribution best fit the expression levels of isoforms in bulk RNA-seq. In our second implemented model, we attempt to infer the probability of each isoform being ‘chosen’ to be expressed in a cell. We calculate the probability of an isoform being chosen based on the observed probability of the isoform being detected. Our third model is identical to the second except that we allow the probability of an isoform being ‘chosen’ to vary between cells. We achieve this by sampling the probability of an isoform being chosen from a Beta distribution, using a similar approach as Velten et al [4]. In our final model, we choose a random number between 0 and 1 for each isoform. The random number is assigned to be that isoform’s probability of being chosen, weighted against the probabilities of the gene’s other isoforms being chosen. For brevity, we will refer to these four models as the Weibull model, the inferred probabilities model, the cell variability model and the random model below.

Figure 3 shows the distributions of the mean number of detected isoforms when one, two, three or four isoforms are expressed per gene per cell for each model. Importantly, the distributions visibly differ between models. To quantitatively confirm this, we perform a K-sample Anderson-Darling test on each row of graphs in Figure 3. We find that the distributions for 1, 2 and 3 isoforms significantly differ between the isoform choice models (p*<*0.001, see Supplementary Tables for details). In contrast, the distributions for 4 isoforms have a p-Value of 1.0, consistent with these distributions originating from the same population. This is as expected, as in the 4 isoform simulations all of the isoforms are picked, and thus we would not expect isoform choice to matter. Our qualitative and quantitative analyses indicate that different mechanisms of isoform choice alter our ability to detect splice isoforms in scRNA-seq. Therefore, a better understanding of the mechanism of isoform choice across the transcriptome could be key to enabling splicing analysis using scRNA-seq data. Without knowing how best to model isoform choice, our results suggest the presence of a substantial confounder.

**Figure 3:**
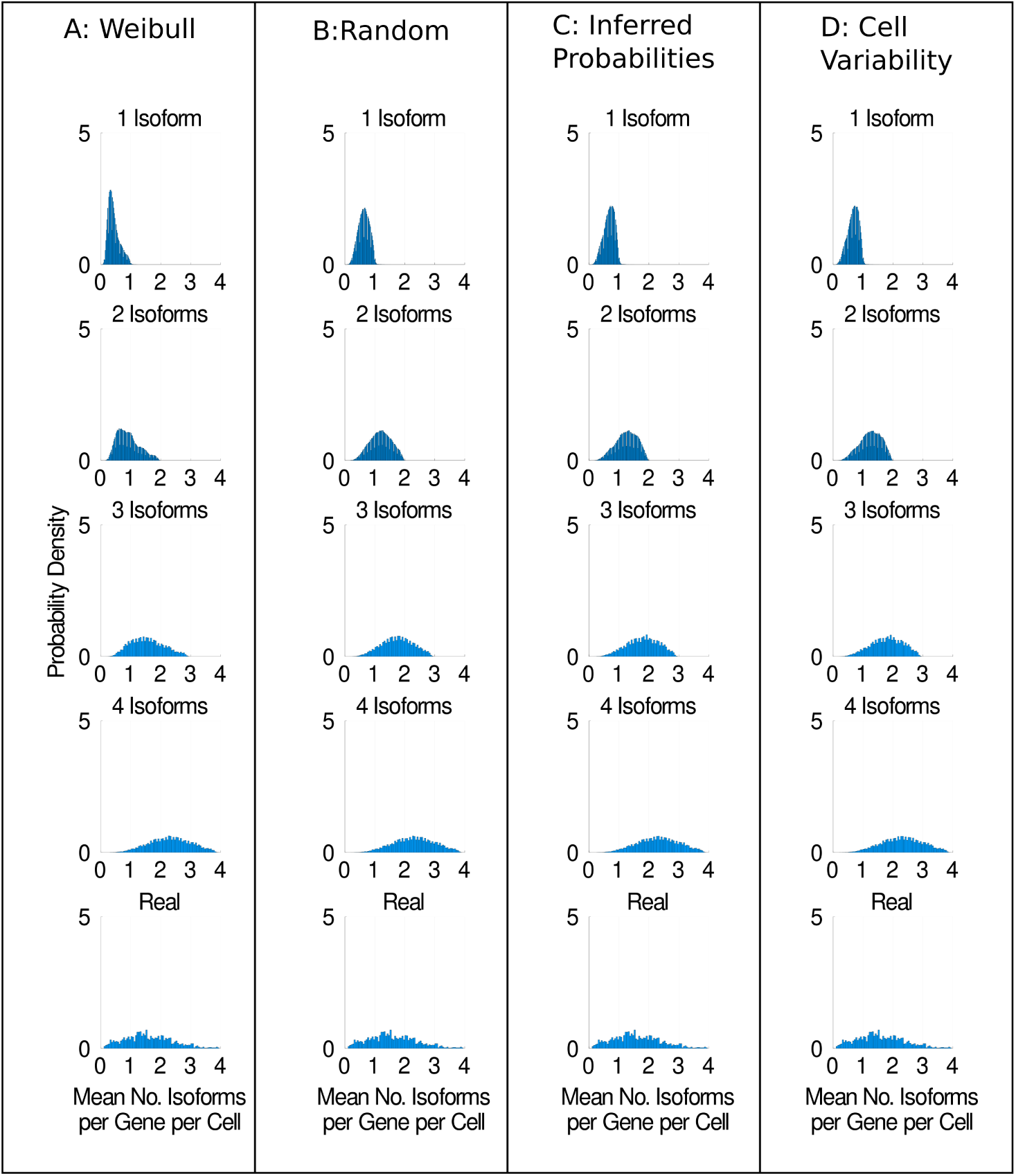
Different models of isoform choice alter our ability to detect isoforms. **a** Distributions of the mean number of isoforms detected per gene per cell for H1 hESCs sequenced at approximately 4 million reads per cell using the Weibull model of isoform choice. **b** shows the same distributions when the random model is used. **c** shows the distributions when the inferred probabilities model is used. **d** shows the distributions when the cell variability model is used. See the main text for a detailed description of each model. Equivalent plots for the other scnorm datasets can be found in Supplementary Figures 2-4.

Interestingly, our simulation results when using the inferred probability model compared with the cell variability model are almost identical. Given that the only difference between these models is whether or not isoform preference is allowed to vary between cells, this indicates that cellular heterogeneity in isoform preference does not change our ability to detect isoforms under the inferred probability model. We perform a K-sample Anderson-Darling test between the inferred probabilities and cell variability models for each row of Figure 3, and we find that these distributions do not significantly differ (see Supplementary Tables). Interestingly, the results of the random model of isoform choice look more like the inferred probability and cell variability models than the Weibull model. This could be because the Weibull model determines the probability of an isoform being chosen based on the rank of that isoform, whereas all of the other models do not use a rank based approach. These observations and the difficulty we have interpreting them illustrate the need for a better understanding of how best to model isoform choice.

### Some models of isoform choice are more plausible than others

In the previous section, we observed that our simulation results for the inferred probability and cell variability models were extremely similar. To investigate how general our observation that allowing isoform preference to vary between cells does not alter our simulation results is, we developed three additional models of isoform choice. In the first model, the probability of selecting each isoform was sampled from a truncated Normal distribution with a mean of 0.25 and a standard deviation of 0.06 in each cell. In the second model, we sample the probability of selecting each isoform from a Bernoulli distribution, in which the value 1 is chosen 25% of the time and the value 0 is chosen 75% of the time in each cell. In the final model, the probability of selecting each isoform is always 0.25 (the ‘p=0.25’ model). The three models are illustrated in Figure 4a and additional details are given in Methods. Under the Normal and the Bernoulli models, the probability of picking each isoform varies between cells, whereas the probability of picking each isoform is constant between cells under the p=0.25 model. Importantly, although the distributions we are sampling isoforms from have very different shapes, the mean probability of picking each isoform is 0.25 for all three distributions.

**Figure 4:**
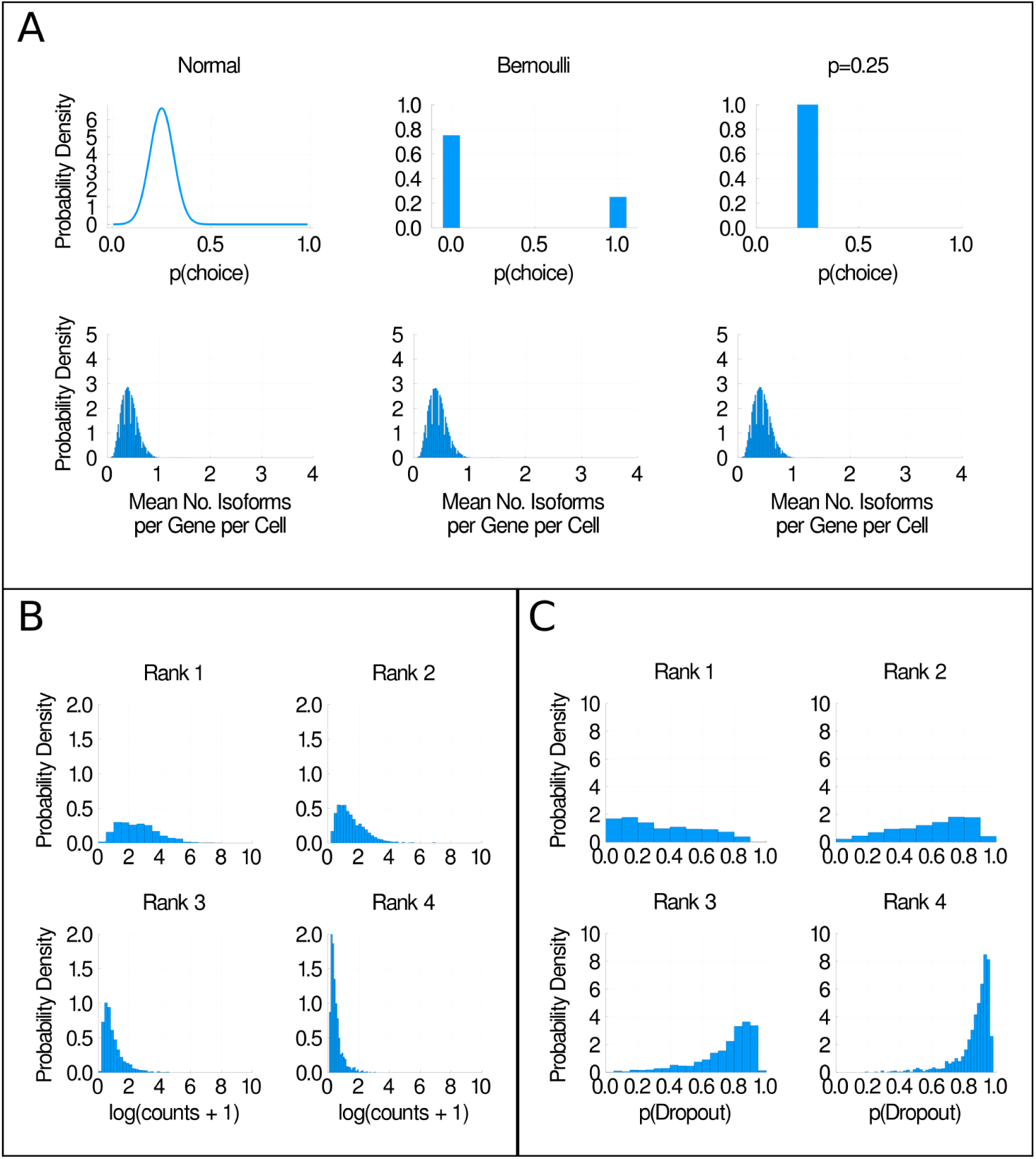
Some models of isoform choice are more plausible than others. **a** We model the probability of picking any given isoform as a Normal distribution, a Bernoulli distribution and a constant probability, all with the same mean (0.25) (top row of graphs). In the bottom row, we show the distributions of the mean number of isoforms per gene per cell detected when each model of isoform choice is used. **b** Histograms of mean isoform expression, ordered by isoform rank. **c** Histograms of dropout probability, ordered by isoform rank. All plots shown are for H1 hESCs sequenced at 1 million reads per cell.

In the second row in Figure 4a, we show the distribution of the mean number of isoforms detected per gene per cell when we simulate one isoform being expressed per gene per cell. There is no visible difference between our simulation results regardless of which model of isoform choice is used. This is supported by a non-significant result in a K-sample Anderson-Darling test (p = 0.998). These findings are consistent with the hypothesis that our simulation results are unchanged whether or not the model of isoform choice used allows cell variability in isoform choice. We suggest that this is because we are reporting the mean number of isoforms detected per gene per cell in our simulations. Across many cells and rounds of simulation, the mean probability of selecting isoforms seems to determine the shape of our simulation result distributions, whereas the higher moments of the isoform choice probability distribution are apparently unimportant. Thus, including cell variability in our isoform choice model appears to not matter. For future scRNA-seq studies in which the mean number of isoforms detected per gene per cell is an important metric, we conjecture that there is no need to model cellular variability in isoform choice, regardless of whether or not such variability exists in reality. Of course, if future studies are interested in precisely what isoforms are present in individual cells rather than a population mean, understanding whether or not cell variability in isoform choice exists is likely to be important.

We have established that our ability to detect isoforms using scRNA-seq is severely affected by the high rate of dropouts in scRNA-seq. Therefore, attempts to infer a biologically meaningful model of isoform choice from scRNA-seq data are likely to fail. However, we can make some general observations to help rule out certain models of isoform choice. In Figure 4b, we have ranked isoforms by their mean expression relative to other isoforms from the same gene (so for example, an isoform with rank 1 has the highest mean expression, an isoform with rank 2 has the second highest mean expression, and so on). Unsurprisingly, we find that the most highly ranked isoforms are substantially more highly expressed than lowly ranked isoforms. This is consistent with the finding that many genes appear to have a ‘major’, more highly expressed isoform, and one or more ‘minor’, less highly expressed isoform [12, 13]. We suggest that this behaviour needs to be represented in some way in future models of isoform choice, and models that do not represent it (for example, our Random, Normal, Bernoulli and p=0.25 models) are probably overly simplistic. In Figure 4c we rank isoforms by their probability of dropout, where the isoform with the lowest probability of dropout compared to other isoforms from the same gene has rank 1. We observe a very similar pattern in which highly ranked isoforms have a substantially lower probability of dropout relative to lowly ranked isoforms, further supporting the finding that ‘major’ and ‘minor’ isoforms exist for many genes.

### A mixture modelling approach suggests genes for which four isoforms are detected typically express around three isoforms per cell

We ask whether our simulation based approach could shed any light on the biological question of how many isoforms are expressed per gene per cell. To do this, we simulate one, two, three and four isoforms being expressed per gene per cell and compare the mean isoforms detected distributions to the distribution of isoforms detected per gene per cell for genes for which four isoforms were detected in the real dataset (see Figure 5a and b). We then approximate each distribution as a log normal distribution and take a mixture modelling approach to estimate the mixing fraction for each of our simulated distributions in the real distribution.

**Figure 5:**
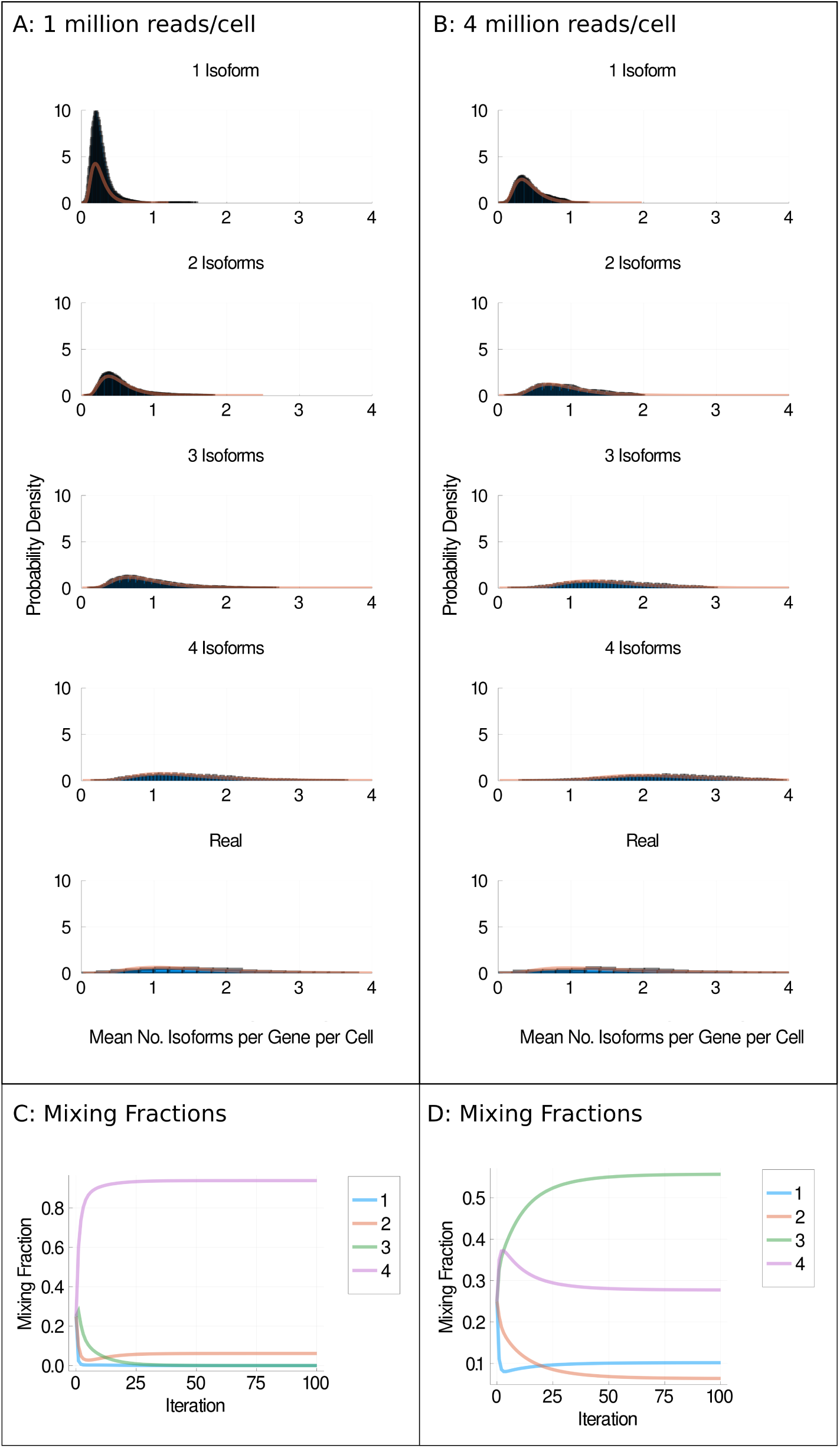
Mixture models. **a** and **b** Distributions of detected isoforms per gene per cell (blue) and log normal fitted distributions (orange) for H1 cells sequenced at 1 million reads per cell (**a**) or 4 million reads per cell (**b**) under the Weibull model. **c** and **d** Mixing fractions vs iterations of expectation maximisation for 1 million reads per cell (**c**) and 4 million reads per cell (**d**). Each coloured line represents the distributions for one, two, three or four isoforms being simulated as expressed per gene per cell. Equivalent plots for other isoform choice models and H9 cells can be found in Supplementary Figures 9-15.

Figure 5c shows the mixing fractions found over 100 iterations of expectation maximisation for H1 hESCs sequenced at approximately 1 million reads per cell. In Figure 5c, the mixing fraction for the distribution corresponding to four isoforms being expressed per gene per cell is over 90%T? his suggests that genes detected to express four isoforms in this dataset typically express four isoforms per gene per cell. However, in Figure 5d, after 100 iterations of expectation maximisation for H1 hESCs sequenced at 4 million reads per cell, the distribution with the largest mixing fraction is that corresponding to three isoforms per gene per cell. This suggests that genes detected to express four isoforms in this dataset most often express three isoforms per gene per cell. As the cDNA sequenced at 1 and 4 million reads per cell came from the same population of cells, it is unlikely that both of these statements are true. We propose several possible explanations for why we might observe this result.

First, we might be over-estimating the dropout rate at 1 million reads per cell. As there is less information with which to infer the dropout rate at 1 million reads per cell compared to at 4 million reads per cell, it is plausible that our estimates of the dropout rate are less accurate at 1 million reads per cell. Whether or not there is a systematic bias towards over-estimating the dropout rate at low sequencing depths is unknown, and goes beyond the scope of this paper.

Second, we have established that the model of isoform choice influences the outcome of our simulations but we do not know which model of isoform choice is correct. Therefore we are (almost certainly) attempting to fit distributions that do not represent reality. Figure 5 shows our mixture modelling approach using the Weibull model of isoform choice. We note however that fitting our alternative models of isoform choice achieves a similar result, in that the largest mixing fraction goes to four isoforms at 1 million reads per cell and to three isoforms at 4 million reads per cell (see Supplementary Figures 9-15).

Third, the genes detected to express four isoforms differ between the sequencing depths of 1 and 4 million reads. More genes are detected to express four isoforms at 4 million reads (1443 versus 1543 for the H1 cells, 1453 versus 1524 for the H9 cells). Whilst this is not a dramatic difference, it does mean that the mixing fractions between these two depths could genuinely differ, although this is unlikely to fully explain the observed difference.

Fourth, we assume all genes for which four isoforms are detected in the real data actually express four isoforms. Due to dropouts and quantification errors, this may not be accurate, and some genes for which four isoforms are detected may express a different number of isoforms in reality.

Fifth, our parameter estimation for quantification errors and isoform choice modelling is not one hundred percent accurate. We can not rule out that this could be confounding the results of our mixture modelling approach.

Our mixture modelling experiments broadly support the hypothesis that it might be common for a cell to produce more than one isoform per gene. However, there are clearly a lot of potential confounders in our approach, many of which relate to uncertainty about dropouts, quantification errors and isoform choice. We note that without having either a ground truth knowledge of how many isoforms are produced from given genes in given cells, or good estimates of dropout probabilities, quantification errors and isoform choice mechanism, it is hard to imagine how an accurate and reliable estimate of the number of isoforms produced per gene per cell could be obtained.

## Discussion

In this study, we use a novel simulation based approach to ask whether it is possible to study alternative splicing at the level of individual cells using scRNA-seq. In our simulations, we simulate four scenarios in which every gene produces one, two, three or four isoforms per gene per cell. That we struggle to clearly distinguish between these four situations emphasises the challenges associated with distinguishing the much subtler and more complex patterns of alternative splicing that likely exist in reality.

We next ask what limitations must be overcome to make alternative splicing analysis possible using scRNA-seq. We find that reducing the probability of dropouts improves our ability to accurately detect isoform number. Therefore, reducing the frequency of dropouts could be one method to improve the accuracy of splicing analyses in scRNA-seq. To some extent, this could be achieved by sequencing cells more deeply, although we note that at 4 million reads per cell we still substantially underestimate isoform number in the H1 hESCs. Unfortunately, extremely deeply sequenced datasets (eg. *>*10 million reads per cell) are likely to suffer more with PCR artefacts and potentially a higher false positive rate of isoform detection [11, 22]. Fundamentally, the low capture efficiency of scRNA-seq is likely a consequence of a small amount of starting material. This can probably be rescued to some extent by more PCR cycles and sequencing at higher depths, however we would not expect this to fully solve the problem.

A more radical way to overcome confounders due to dropouts would be if scRNA-seq technologies changed in some fundamental way that increased capture efficiency. Whether this is feasible is unclear. Alternatively, we note that if we could estimate the probability of dropout for each isoform more accurately, in theory it should be possible to correct for confounding due to dropouts in splicing analyses. Therefore, to enable splicing analysis using scRNA-seq, either the capture efficiency of the technology needs to improve, or more work characterising the probability of dropouts at an isoform level is required.

Long read technologies could in theory enable 100% accurate isoform quantification, if issues due to a high base calling error rate could be overcome [23]. However we find that even when no isoform detection errors occur, our ability to accurately detect isoforms is very limited. Therefore, long read technologies or isoform quantification software improvements alone are not sufficient to enable accurate splicing analysis in scRNA-seq.

Little is known about the biological process of isoform choice in individual cells for most genes. Thus, accurately modelling this process is challenging. We find that different models of isoform choice alter our simulation results. This indicates that without better understanding of the process of isoform choice, alternative splicing analyses are potentially confounded by this unknown factor. Research into the process of isoform choice within individual cells across the transcriptome would enable more accurate models of isoform choice to be built, reducing or removing this confounder from future alternative splicing analyses. We observe that when studying the mean number of isoforms detected per gene per cell, it appears to be unimportant whether or not there is cell variability in isoform choice from a modelling perspective. Of course, if the goal is to accurately detect which isoforms are present in each cell, establishing whether cell variability exists and modelling any variability will be essential. However, we note that imputation remains challenging and often inaccurate at the gene level for scRNA-seq [24]. We therefore anticipate it will be some time before accurate imputation is feasible at the isoform level.

We are able to detect evidence in support of ‘major’ and ‘minor’ isoforms, and propose that future models of isoform choice should attempt to capture this behaviour. However, we note that whilst our observations help discard models of isoform choice, we believe that scRNA-seq is currently too confounded by dropouts to accurately infer a model of isoform choice at the single cell level. We suggest that smFISH would be a more appropriate technology to investigate how isoform choice is regulated in individual cells. Indeed, smFISH has previously been used to study alternative splicing and isoform choice in individuals cells for a small number of genes [4, 3, 6]

The results of our mixture modelling experiments are consistent with multiple isoforms being produced per gene per cell, however we note that our mixture modelling experiments are heavily confounded by a lack of understanding about dropouts, isoform choice and perhaps quantification errors to a lesser extent. Therefore, we argue that at this time, scRNA-seq will not be able to provide the answer to basic biological questions about how many isoforms are produced per gene per cell.

Based on our findings, at this time we do not recommend attempting alternative splicing analysis using scRNA-seq. However, we make actionable suggestions for how splicing analysis could be enabled in the future. An improved understanding of the prevalence of technical dropouts at the isoform level could enable us to reduce confounding due to dropouts. Improvements to the capture efficiency of scRNA-seq would similarly reduce confounding. Increased study of isoform choice at the single cell level using technologies such as smFISH would enable better models of isoform choice to be generated, eliminating confounders. Although we find quantification errors to be a relatively small confounder, further reducing quantification errors using long read technologies and more accurate quantification tools would be welcome. Although we have concluded that accurate alternative splicing analysis with scRNA-seq is not possible today, we are optimistic that it could become possible in the near future.

## Conclusions

At present, alternative splicing analyses using scRNA-seq are substantially confounded. Better characterisation of dropouts or improvements in capture efficiency would reduce confounding due to dropouts. Further research into the process of isoform choice at a single cell level would reduce confounding due to a lack of knowledge about isoform choice. Quantification errors are a relatively minor confounder, although improvements in this area are still welcome. At present, to the best of our knowledge, a large scale unconfounded analysis of the number of isoforms produced per gene per cell has not been performed. Therefore, we still do not know how many isoforms are typically produced per gene per cell.

## Methods

### Data Preprocessing

Our simulation approach requires an isoform-cell counts matrix as input. To generate isoform-cell counts matrices, we used Kallisto to quantify reads from each cell against the Gencode mouse vM20 transcriptome for our B lymphocyte and mESC datasets, and against the Gencode human v20 transcriptome for our hESC dataset [25, 26].

### Simulation Approach

Our simulation approach is summarised as an algorithm below.

Step 1: Select genes for which four isoforms are detected in scRNA-seq data

**Algorithm 1:**
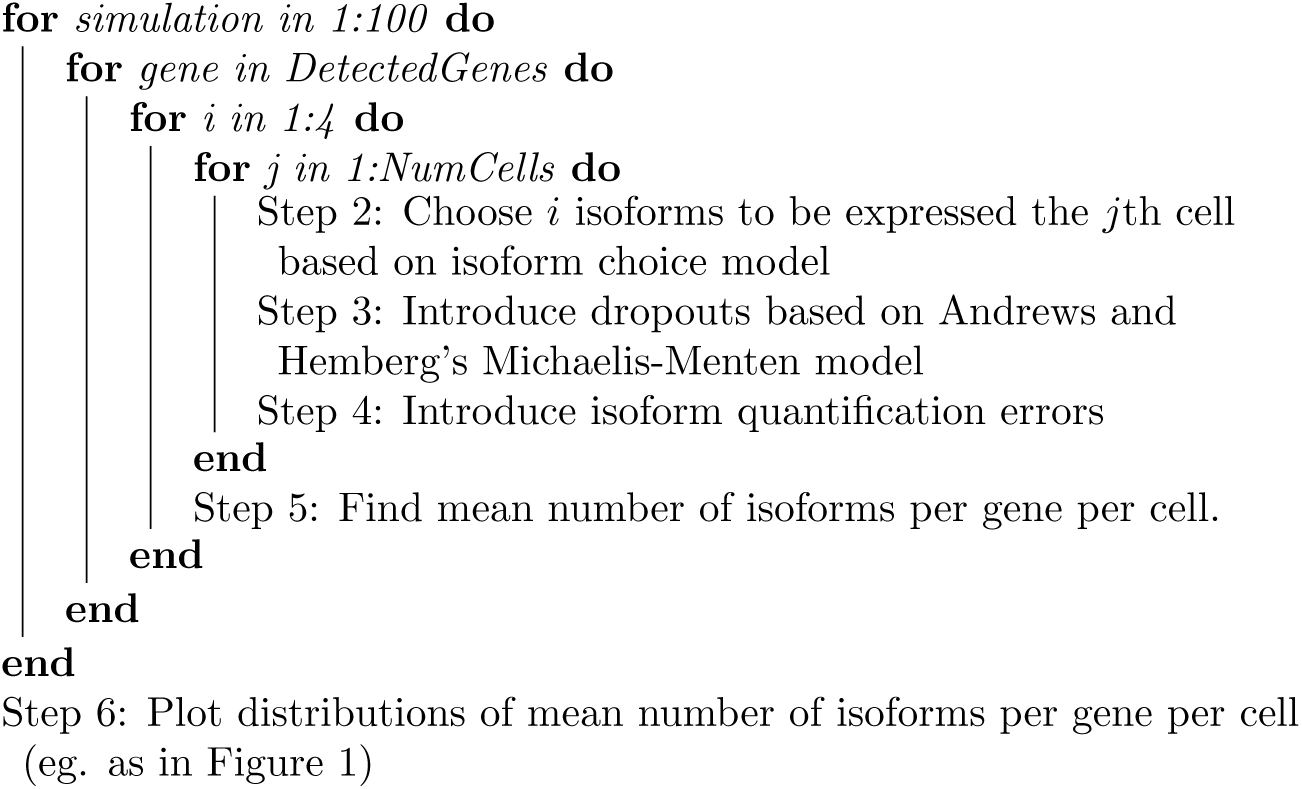
Our simulation approach

We expand upon each step below.

#### Step 1: Select genes for which four isoforms are detected in scRNA-seq data

Our simulation approach takes an isoform-cell counts table as input. We define an isoform as detected if it has more than five counts in at least two cells. We select genes for which exactly four isoforms pass this threshold.

#### Step 2: Choose *i* isoforms to be expressed the *j*th cell based on isoform choice model

In this step, we probabilistically choose *i* isoforms to be expressed in each cell, where *i* is one, two, three or four. The default model used in this study was the Weibull model, which was used to produce all of our main Figures unless otherwise stated.

#### The Weibull model

In [18], Hu et al found that the median frequency, *mf* (*k, M*), of the *k*th dominant isoform of a gene with *M* detected isoforms can be described as:

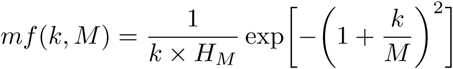

where *H*_*M*_ is the *M* th generalised harmonic number:

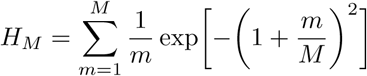

In our implementation of this model of isoform choice, we first rank the isoforms in order of magnitude expression for each gene, with the most highly expressed isoform having rank 1, the second most highly expressed isoform having rank 2 and so on. We calculate magnitude of expression by summing the total number of counts across all cells for that isoform. We then use the median frequency formula above to find the predicted median frequency for each isoform. We define the probability of picking an isoform with rank *k* for a gene with *M* detected isoforms as:

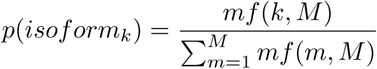

With *M* = 4, the probabilities become [0.55, 0.28, 0.12, 0.05].

#### The inferred probabilities model

In this model, we attempt to infer the probability of an isoform being chosen from its probability of being detected. The formula below relates the probability of choosing an isoform, *P* (*Choice*), to its probability of being detected, *P* (*Detected*):

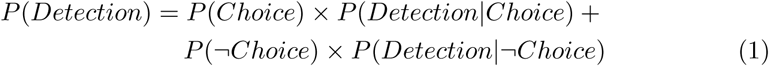

Where *P* (*¬Choice*) is the probability of not choosing an isoform. In practice:

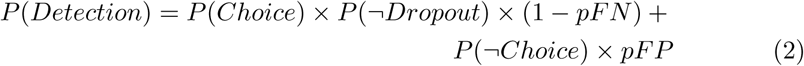

Where *P* (*¬Dropout*) is the probability that there is not a dropout, *pFN* is the probability that there is a false negative event due to a quantification error and *pFP* is the probability that there is a false positive event due to a quantification error. This rearranges to:

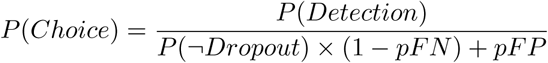

In practice, we sometimes find *P* (*Choice*) is greater than 1, probably because our estimation of *P* (*/Dropout*), *pFN* and/or *pFP* is inaccurate for that isoform. When this occurs, we set *P* (*Choice*) equal to one.

In our simulations, we calculate *P* (*Choice*) for each isoform from a given gene. The probability of picking a particular isoform to be expressed in our simulation is that isoform’s *P* (*Choice*) divided by the sum of *P* (*Choice*)s for that gene’s isoforms.

#### The cell variability model

The cell variability model is identical to the inferred probabilities model except that the probability of picking a given isoform, *i* is allowed to vary between cells. This is acheived by sampling the probability of picking isoform *i* in a given cell *c, p*_*i*_*c*, from a Beta distribution, taking a similar approach to that described in [4] :

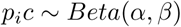

Where

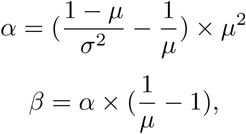

where *µ* is the mean probability of choosing *i* across all cells, i.e. *µ* = *P* (*Choice*). Based on attempts to characterise the mean-variance relationship for the probability of choosing a particular gene by Velten et al [4], we estimate that the sample standard deviation, *σ*, is approximately 0.002. We find *p*_*i*_*c* for each isoform for a given gene. In our simulations, the probability of picking isoform *i* in cell *c* is that isoform’s *p*_*i*_*c* divided by the sum of *p*_*i*_*c*s for that gene’s isoforms.

#### The random model

For this model, each isoform is associated a weight randomly sampled between zero and one. The probability of picking a particular isoform to be expressed in our simulation is that isoform’s weight divided by the sum of all the weights for that gene’s isoforms.

#### The Normal model

The weights for each isoform were sampled from a truncated Normal distribution with a mean of 0.25 and a standard deviation of 0.06. This sampling was performed for each isoform in each cell. Within each cell, the probability of picking a particular isoform to be expressed in our simulation is that isoform’s weight divided by the sum of all the weights for that gene’s isoforms.

#### The Bernoulli model

The weights for each isoform were sampled from a Bernoulli distribution with a mean of 0.25. This sampling was performed for each isoform in each cell. Within each cell, the probability of picking a particular isoform to be expressed in our simulation is that isoform’s weight divided by the sum of all the weights for that gene’s isoforms. If all four isoforms for a given gene had a zero weight, we set the probability of picking each isoform to 0.25.

#### The p=0.25 model

The probability of choosing each isoform was always 0.25.

#### Step 3: Introduce dropouts based on Andrews et al’s Michaelis-Menten model

We calculate the probability of dropouts for each isoform using a Michaelis-Menten model proposed by Andrews and Hemberg [9]. We calculate the probability of dropouts for each isoform as:

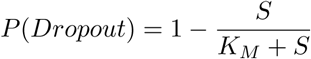

Where *S* is the mean expression of that isoform across cells and *K*_*M*_ is the Michaelis-Menten constant. We estimate the value of *K*_*M*_ for each dataset by applying maximum-likelihood estimation using the equation above and the rate of dropouts and the mean expression of isoforms across the entire transcriptome.

#### Step 4: Introduce quantification errors

Based on our previous benchmarking study [8], we estimate that the probability of a false positive given an isoform has no reads mapping to it, *pFP*, is about and the probability of a false negative given an isoform has reads mapping to it, *pFN*, is about 0.04 for Kallisto when run on full length coverage scRNA-seq data. Unless otherwise stated in the text, these were the error rates applied in our simulations.

#### Step 5: Find mean number of isoforms per gene per cell

After iterating over every cell in our simulation, we sum the the number of isoforms detected in each cell and divide by the number of cells to find the mean number of detected isoforms per gene per cell.

#### Step 6: Plot distributions of mean number of isoforms per gene per cell

Step 5 is carried out in each simulation, for every gene in which four isoforms were detected in the real scRNA-seq data. Consequently a large list of mean number of detected isoforms per gene per cell is generated which we plot as distributions (eg. see Figure 1).

### Mixture Modelling

In our mixture modelling experiments, we begin by fitting log normal distributions to each of our simulation distributions and to the distribution of mean isoforms detected for genes with four detected isoforms in the real data. We then use expectation maximisation to estimate the mixing fraction of each of the simulated distributions in the real distribution. In our expectation step, we calculate the probability that each data point belongs to a given distribution, which we refer to as the responsibility. The responsibility for the *i*th mean number of isoforms per gene per cell and the *c*th simulation distribution is:

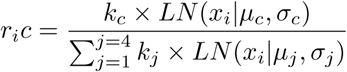

where *k* is the mixing fraction, *x*_*i*_ is the *i*th mean number of isoforms per gene per cell and *LN* (*x*_*i*_|*µ*_*c*_, *σ*_*c*_) is the probability density function for the log normal with mean *µ*_*c*_ and variance 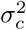. The maximisation function for the mixing fraction is:

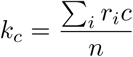

Where *n* is the number of datapoints in *r*_*i*_*c*. Note that we only perform expectation maximisation for the mixing fractions of the distributions and not for the means or standard deviations.

## Competing interests

The authors declare that they have no competing interests.

### Author’s contributions

JW, AFS and MH conceived the study and designed the experiments. JW and PA carried out the experiments. JW wrote the manuscript. JW, AFS and MH supervised the experiments.

#### Acknowledgements

We would like to thank Stephanie Telerman for helpful discussions.

## Availability of data and materials

The Kolodziejczyk et al. mESC data was accessed from the ArrayExpress database (http://www.ebi.ac.uk/arrayexpress) using the accession number E-MTAB-2600, as described in the Kolodziejczyk et al. paper [27]. The BLUEPRINT data was accessed under GEO accession number GSE106663 [28]. The hESC datasets were accessed under GEO accession number GSE85917 [19]. Our quantification pipelines, which download scRNA-seq data, perform transcript level quantification and generate an isoform-cell matrix, can be found at: https://github.com/jenni-westoby/Isoform_Cell_Matrix_Generation. Our simulation pipeline can be found at: https://github.com/jenni-westoby/Obstacles.

## Figures

### Additional Files

#### Additional file 1 — Supplementary Figures and Tables

This PDF file contains all of the supplementary figures and tables for this paper.

