## Supplementary Figures and Tables for "Obstacles to Studying Alternative Splicing Using scRNA-seq"

October 4, 2019

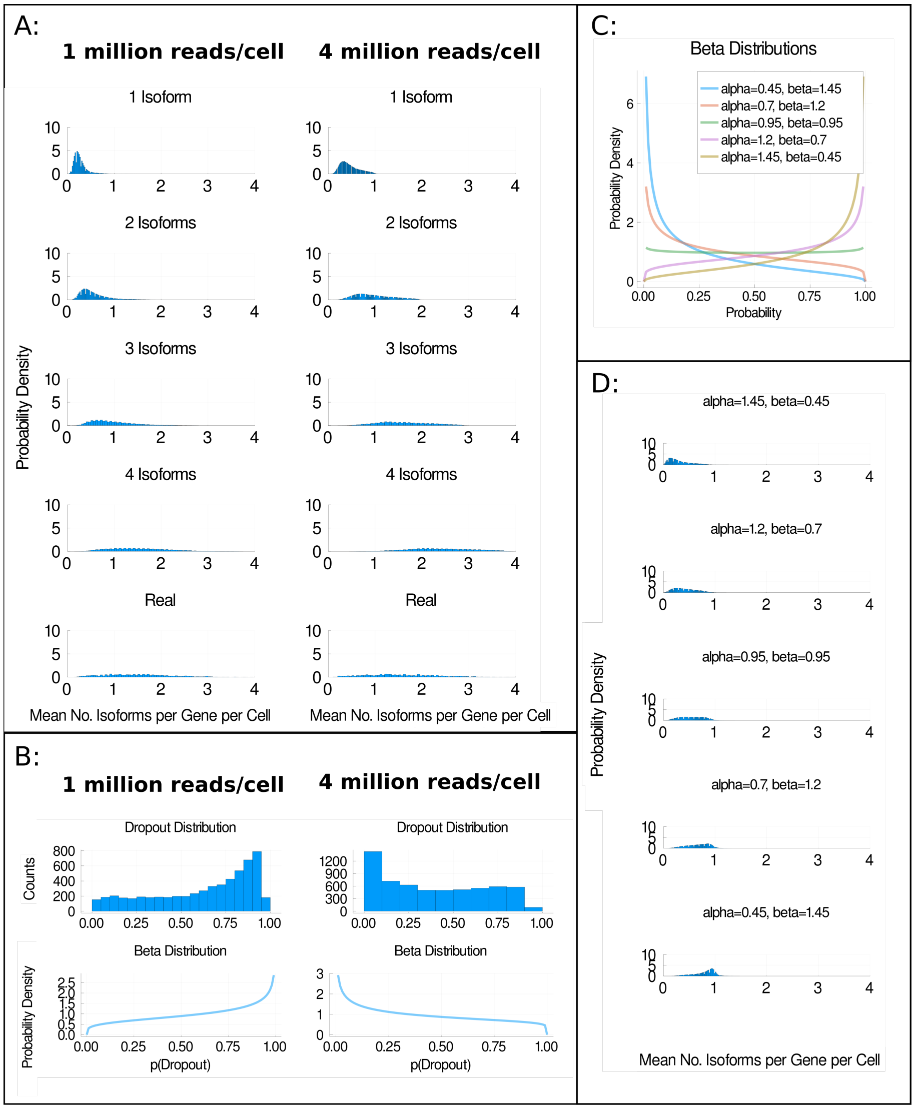


Figure 1: The impact of dropouts on isoform detection. **a** Distributions of the mean number of isoforms detected per gene per cell for H9 hESCs whose cDNA was split and sequenced at approximately 1 million reads per cell or 4 million reads per cell on average [1]. **b** shows the distribution of the probabilities of dropouts (p(Dropout)) in each group and an approximation of these distributions using a Beta distribution. **c** shows five Beta Distributions from which dropout probabilities were sampled from in the simulations used to generate **d**. In **d**, the distribution of the mean number of isoforms detected per gene per cell is shown for simulations in which one isoform was produced per gene per cell. Each plot corresponds to a simulation in which dropout probabilities were sampled from one of the distributions shown in **c**.

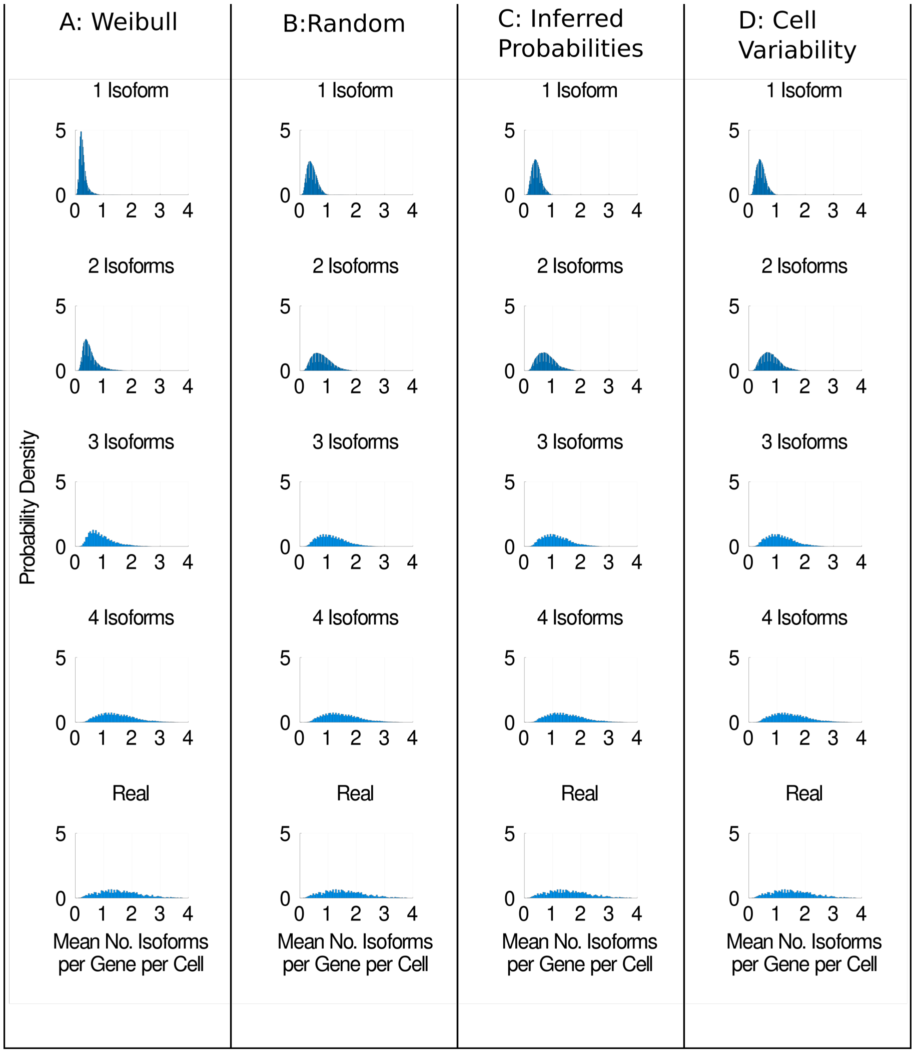


Figure 2: Different models of isoform choice alter our ability to detect isoforms. **a** Distributions of the mean number of isoforms detected per gene per cell for H1 hESCs sequenced at approximately 1 million reads per cell using the Weibull model of isoform choice [1, 2]. **b** shows the same distributions when the random model is used. **c** shows the distributions when the inferred probabilities model is used. **d** shows the distributions when the cell variability model is used. See the main text for a detailed description of each model.

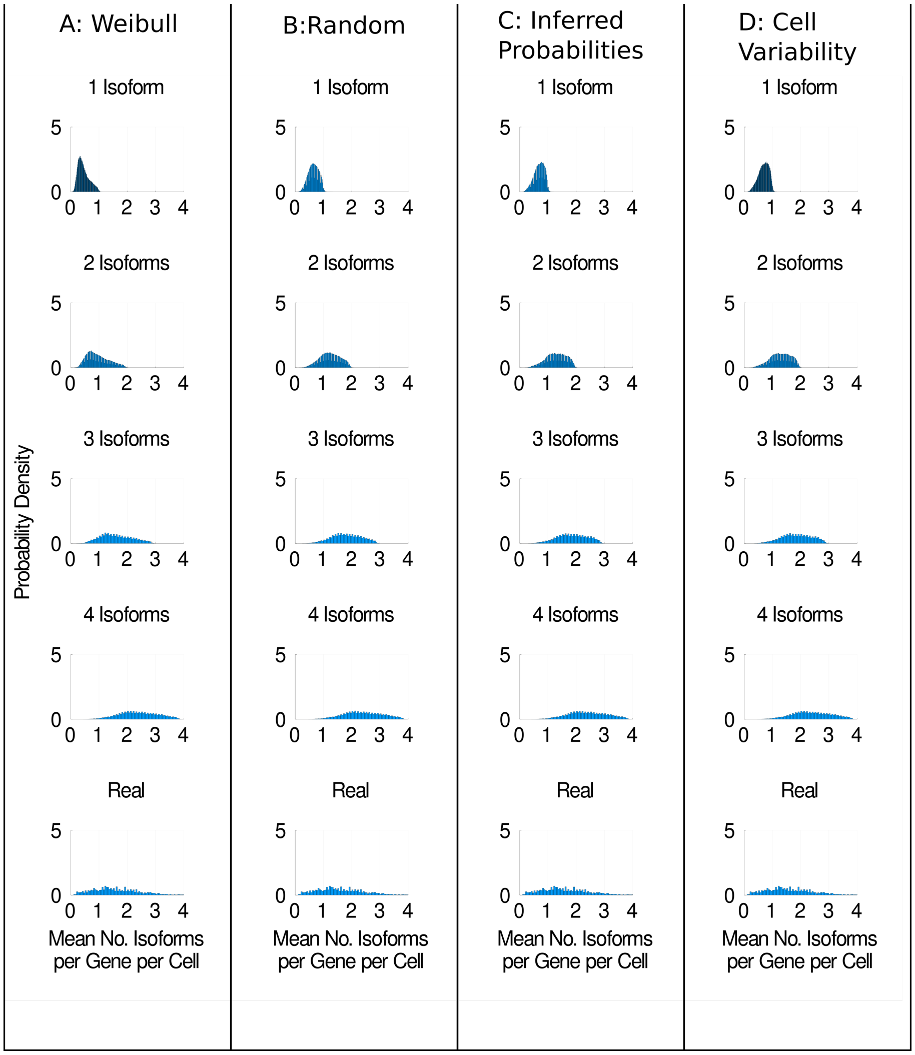


Figure 3: Different models of isoform choice alter our ability to detect isoforms. **a** Distributions of the mean number of isoforms detected per gene per cell for H9 hESCs sequenced at approximately 4 million reads per cell using the Weibull model of isoform choice [1, 2]. **b** shows the same distributions when the random model is used. **c** shows the distributions when the inferred probabilities model is used. **d** shows the distributions when the cell variability model is used. See the main text for a detailed description of each model.

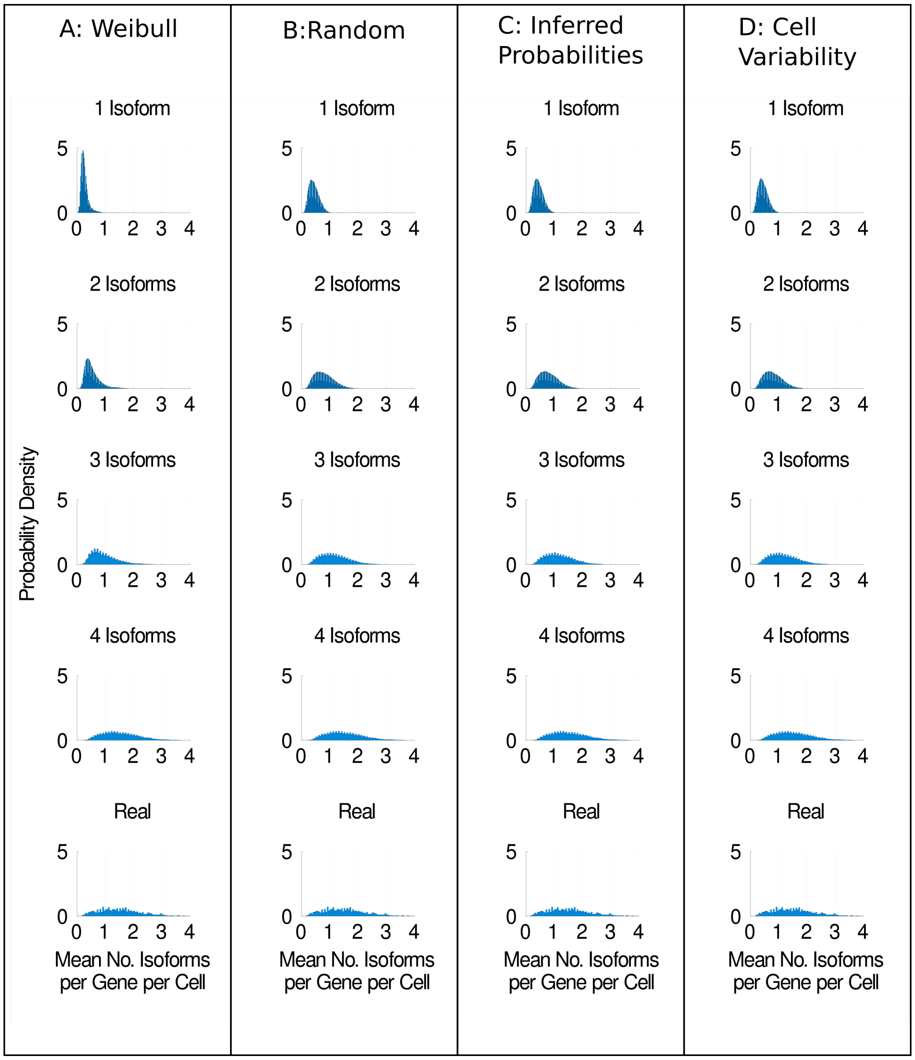


Figure 4: Different models of isoform choice alter our ability to detect isoforms. **a** Distributions of the mean number of isoforms detected per gene per cell for H9 hESCs sequenced at approximately 1 million reads per cell using the Weibull model of isoform choice [1, 2]. **b** shows the same distributions when the random model is used. **c** shows the distributions when the inferred probabilities model is used. **d** shows the distributions when the cell variability model is used. See the main text for a detailed description of each model.

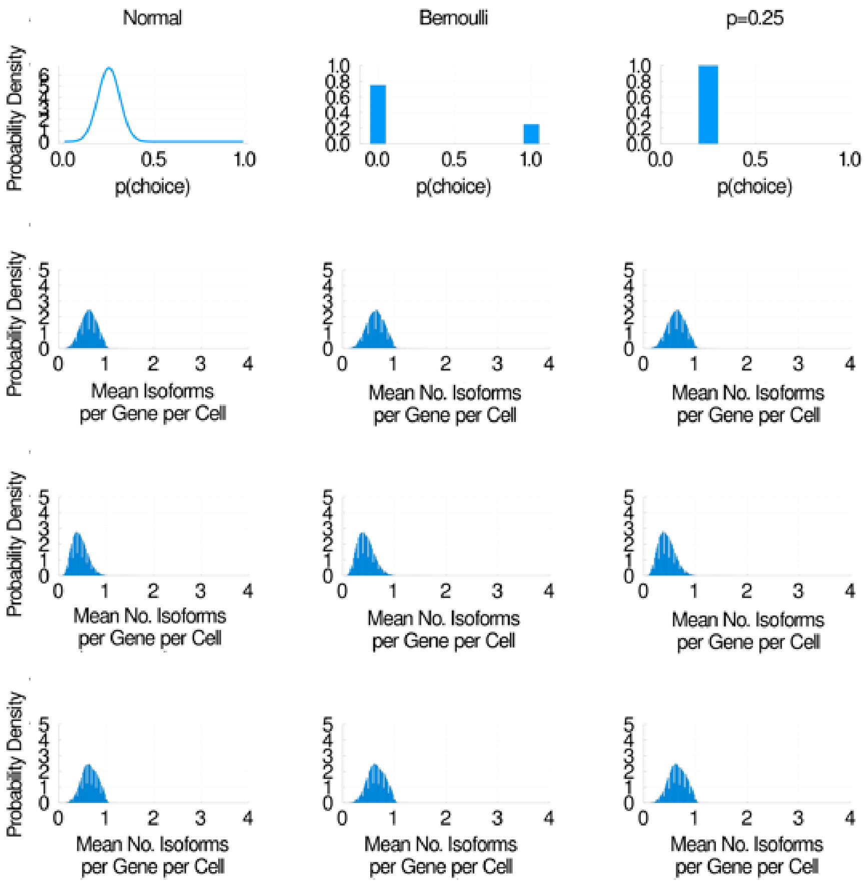


Figure 5: Some models of isoform choice are more plausible than others. We model the probability of picking any given isoform as a Normal distribution, a Bernoulli distribution and a constant probability, all with the same mean (0.25) (top row of graphs). In the following rows, we show the distributions of the mean number of isoforms per gene per cell detected when each model of isoform choice is used. The second row is H1 hESCs sequenced at 4 million reads, the third row is H9 hESCs sequenced at 1 million reads, the fourth row is H9 hESCs sequenced at 4 million reads.

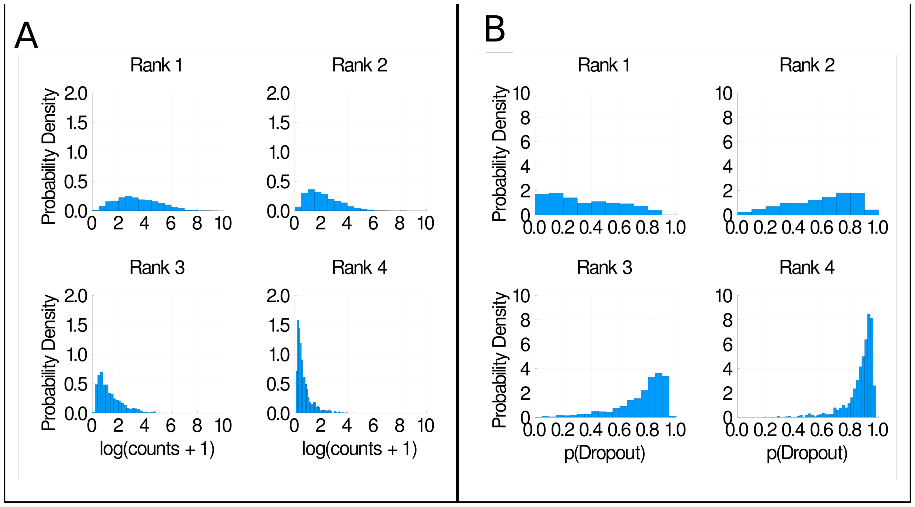


Figure 6: **a** Histograms of mean isoform expression, ordered by isoform rank. **b** Histograms of dropout probability, ordered by isoform rank. All plots shown are for H1 hESCs sequenced at 4 million reads per cell.

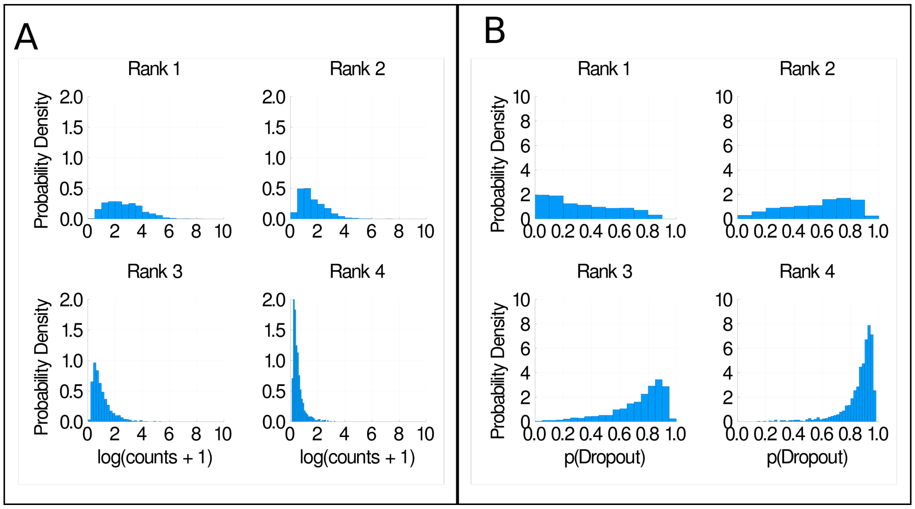


Figure 7: **a** Histograms of mean isoform expression, ordered by isoform rank. **b** Histograms of dropout probability, ordered by isoform rank. All plots shown are for H9 hESCs sequenced at 1 million reads per cell.

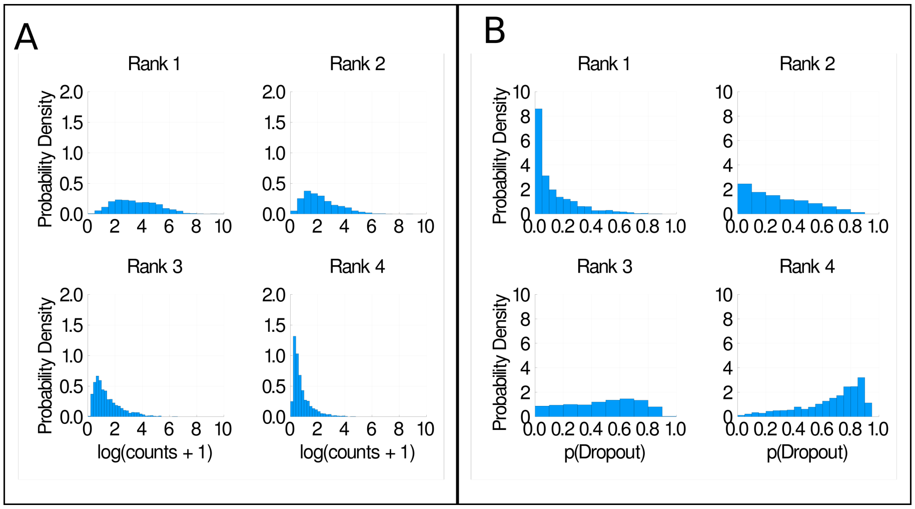


Figure 8: **a** Histograms of mean isoform expression, ordered by isoform rank. **b** Histograms of dropout probability, ordered by isoform rank. All plots shown are for H9 hESCs sequenced at 4 million reads per cell.

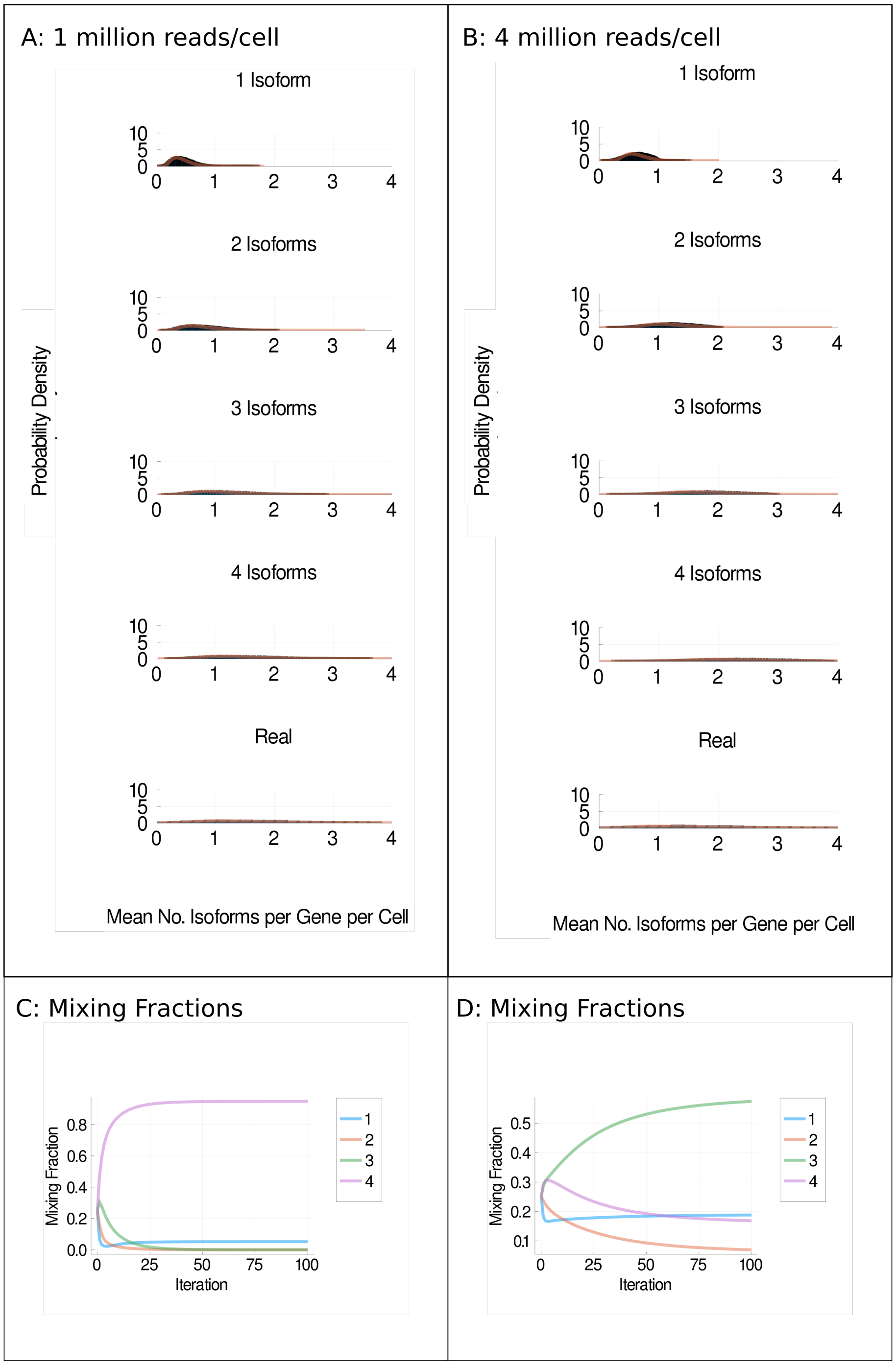


Figure 9: Mixture models. **a** and **b** Distributions of detected isoforms per gene per cell (blue) and log normal fitted distributions (orange) for H1 cells sequenced at 1 million reads per cell (**a**) or 4 million reads per cell (**b**) under the random model [1]. **c** and **d** Mixing fractions vs iterations of expectation maximisation for 1 million reads per cell (**c**) and 4 million reads per cell (**d**). Each coloured line represents the distributions for one, two, three or four isoforms being simulated as expressed per gene per cell.

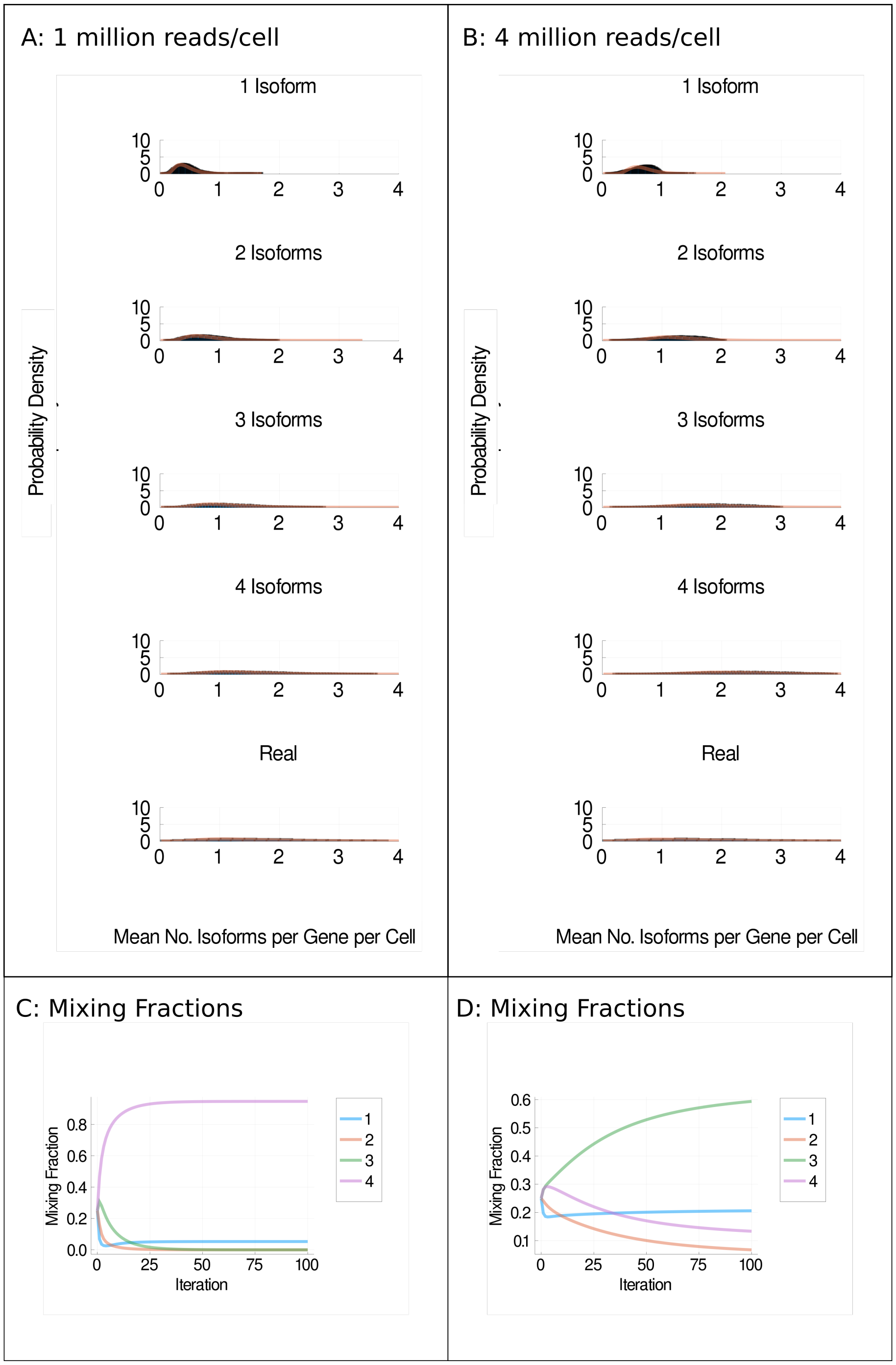


Figure 10: Mixture models. **a** and **b** Distributions of detected isoforms per gene per cell (blue) and log normal fitted distributions (orange) for H1 cells sequenced at 1 million reads per cell (**a**) or 4 million reads per cell (**b**) under the inferred model [1]. **c** and **d** Mixing fractions vs iterations of expectation maximisation for 1 million reads per cell (**c**) and 4 million reads per cell (**d**). Each coloured line represents the distributions for one, two, three or four isoforms being simulated as expressed per gene per cell.

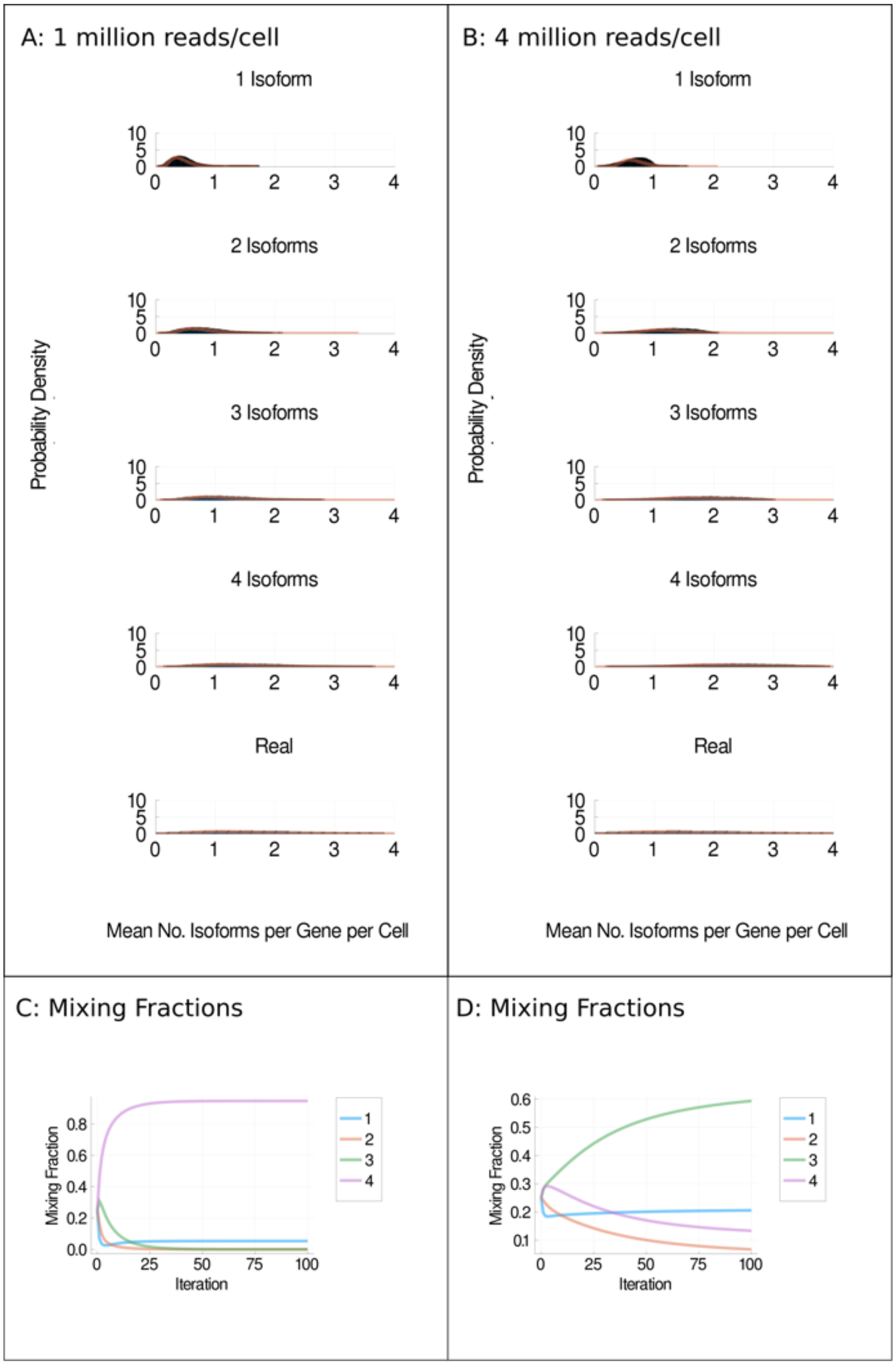


Figure 11: Mixture models. **a** and **b** Distributions of detected isoforms per gene per cell (blue) and log normal fitted distributions (orange) for H1 cells sequenced at 1 million reads per cell (**a**) or 4 million reads per cell (**b**) under the cell variable model [1, 3]. **c** and **d** Mixing fractions vs iterations of expectation maximisation for 1 million reads per cell (**c**) and 4 million reads per cell (**d**). Each coloured line represents the distributions for one, two, three or four isoforms being simulated as expressed per gene per cell.

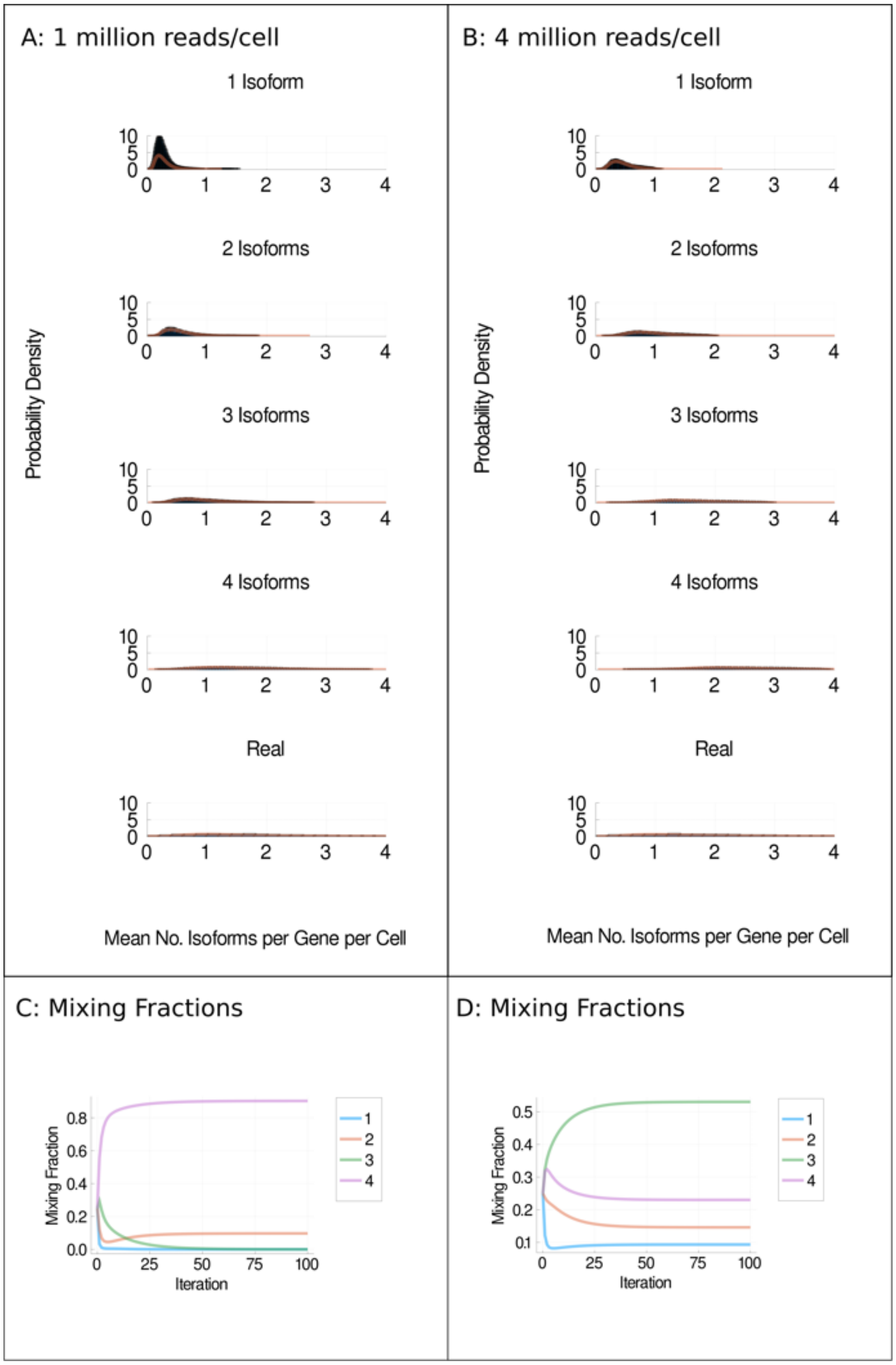


Figure 12: Mixture models. **a** and **b** Distributions of detected isoforms per gene per cell (blue) and log normal fitted distributions (orange) for H9 cells sequenced at 1 million reads per cell (**a**) or 4 million reads per cell (**b**) under the Weibull model [1, 2]. **c** and **d** Mixing fractions vs iterations of expectation maximisation for 1 million reads per cell (**c**) and 4 million reads per cell (**d**). Each coloured line represents the distributions for one, two, three or four isoforms being simulated as expressed per gene per cell.

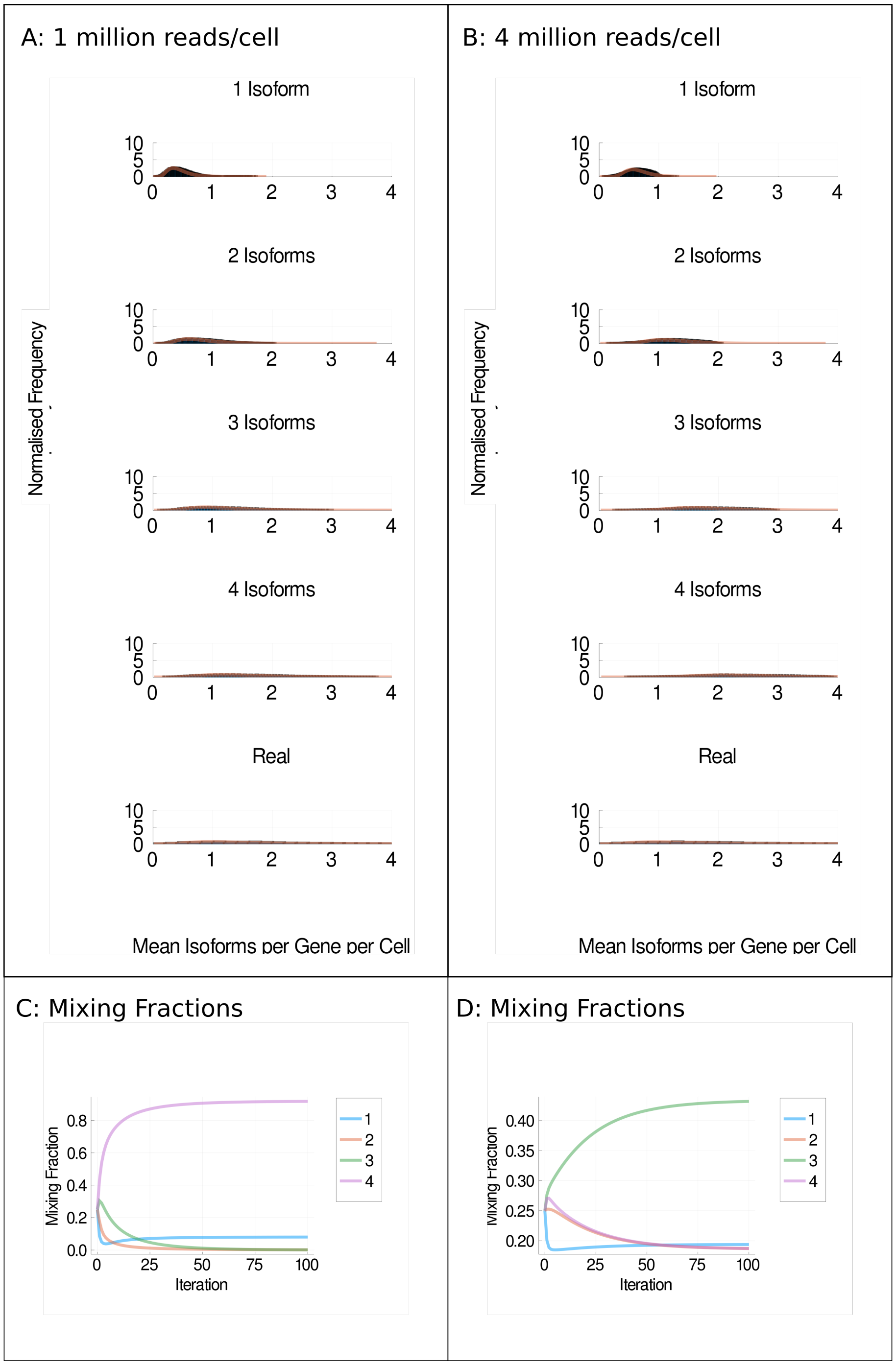


Figure 13: Mixture models. **a** and **b** Distributions of detected isoforms per gene per cell (blue) and log normal fitted distributions (orange) for H9 cells sequenced at 1 million reads per cell (**a**) or 4 million reads per cell (**b**) under the random model [1]. **c** and **d** Mixing fractions vs iterations of expectation maximisation for 1 million reads per cell (**c**) and 4 million reads per cell (**d**). Each coloured line represents the distributions for one, two, three or four isoforms being simulated as expressed per gene per cell.

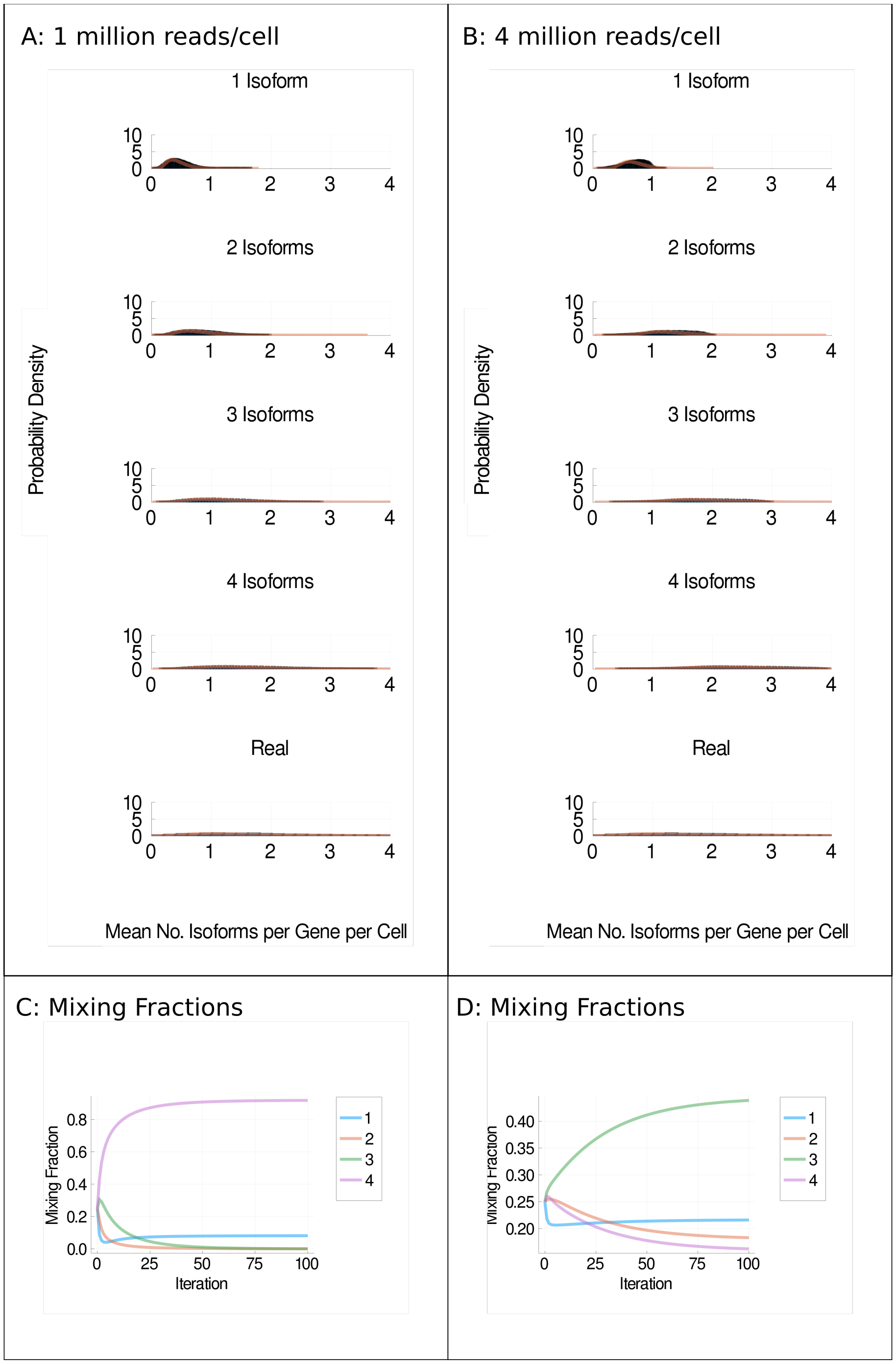


Figure 14: Mixture models. **a** and **b** Distributions of detected isoforms per gene per cell (blue) and log normal fitted distributions (orange) for H9 cells sequenced at 1 million reads per cell (**a**) or 4 million reads per cell (**b**) under the inferred model [1]. **c** and **d** Mixing fractions vs iterations of expectation maximisation for 1 million reads per cell (**c**) and 4 million reads per cell (**d**). Each coloured line represents the distributions for one, two, three or four isoforms being simulated as expressed per gene per cell.

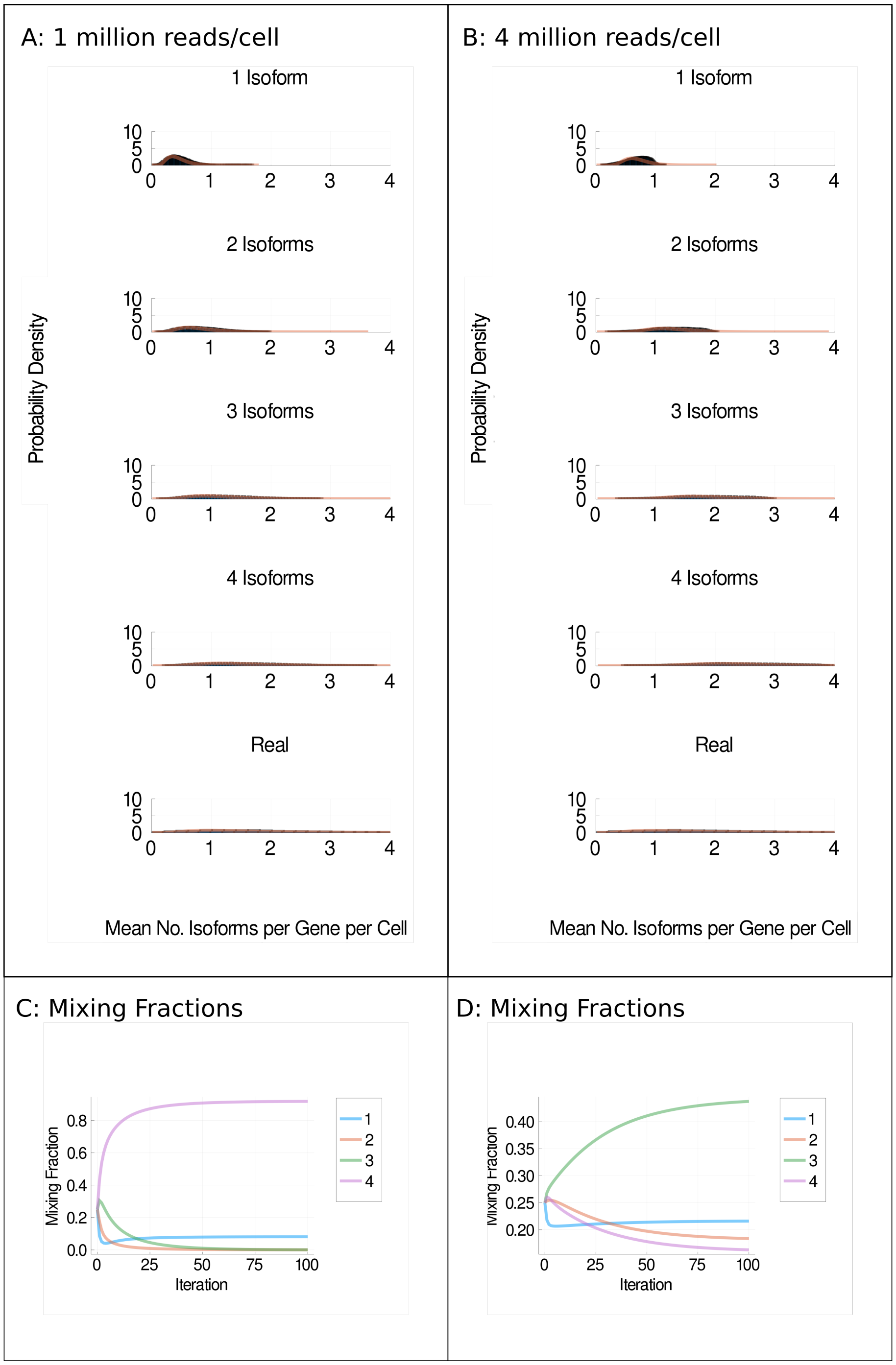


Figure 15: Mixture models. **a** and **b** Distributions of detected isoforms per gene per cell (blue) and log normal fitted distributions (orange) for H9 cells sequenced at 1 million reads per cell (**a**) or 4 million reads per cell (**b**) under the cell variable model [1, 3]. **c** and **d** Mixing fractions vs iterations of expectation maximisation for 1 million reads per cell (**c**) and 4 million reads per cell (**d**). Each coloured line represents the distributions for one, two, three or four isoforms being simulated as expressed per gene per cell.

| No. Isoforms Simulated | p-Value |
|---|---|
| 1 | 0.0 |
| 2 | 0.0 |
| 3 | 0.0 |
| 4 | 1.0 |

Table 1: Results of K-sample Anderson–Darling test, which tests whether multiple collections come from the same population. The test was applied to each row of graphs in Figure 3, in other words testing whether the distributions generated by different isoform choice models are significantly different.

| No. Isoforms Simulated | p-Value |
|---|---|
| 1 | 0.984627 |
| 2 | 0.999594 |
| 3 | 0.998207 |
| 4 | 0.99988 |

Table 2: Results of K-sample Anderson–Darling test, which tests whether multiple collections come from the same population. The test was applied to the simulation results generated using the Inferred Probabilities vs the Cell Variable models of isoform choice in Figure 3 to test whether the distributions generated by different isoform choice models significantly differ.

| No. Isoforms Simulated | p-Value |
|---|---|
| 1 | 0.0 |
| 2 | 0.0 |
| 3 | 0.0 |
| 4 | 0.999994 |

Table 3: Results of K-sample Anderson–Darling test, which tests whether multiple collections come from the same population. The test was applied to each row of graphs in Supplementary Figure 2, in other words testing whether the distributions generated by different isoform choice models are significantly different.

| No. Isoforms Simulated | p-Value |
|---|---|
| 1 | 0.995216 |
| 2 | 0.901331 |
| 3 | 0.961482 |
| 4 | 0.991821 |

Table 4: Results of K-sample Anderson–Darling test, which tests whether multiple collections come from the same population. The test was applied to the simulation results generated using the Inferred Probabilities vs the Cell Variable models of isoform choice in Supplementary Figure 2 to test whether the distributions generated by different isoform choice models significantly differ.

| No. Isoforms Simulated | p-Value |
|---|---|
| 1 | 0.0 |
| 2 | 0.0 |
| 3 | 0.0 |
| 4 | 1.0 |

Table 5: Results of K-sample Anderson–Darling test, which tests whether multiple collections come from the same population. The test was applied to each row of graphs in Supplementary Figure 3, in other words testing whether the distributions generated by different isoform choice models are significantly different.

| No. Isoforms Simulated | p-Value |
|---|---|
| 1 | 0.954213 |
| 2 | 0.99939 |
| 3 | 0.998889 |
| 4 | 0.998885 |

Table 6: Results of K-sample Anderson–Darling test, which tests whether multiple collections come from the same population. The test was applied to the simulation results generated using the Inferred Probabilities vs the Cell Variable models of isoform choice in Supplementary Figure 3 to test whether the distributions generated by different isoform choice models significantly differ.

| No. Isoforms Simulated | p-Value |
|---|---|
| 1 | 0.0 |
| 2 | 0.0 |
| 3 | 0.0 |
| 4 | 0.999993 |

Table 7: Results of K-sample Anderson–Darling test, which tests whether multiple collections come from the same population. The test was applied to each row of graphs in Supplementary Figure 4, in other words testing whether the distributions generated by different isoform choice models are significantly different.

| No. Isoforms Simulated | p-Value |
|---|---|
| 1 | 0.988069 |
| 2 | 0.985811 |
| 3 | 0.999795 |
| 4 | 0.971993 |

Table 8: Results of K-sample Anderson–Darling test, which tests whether multiple collections come from the same population. The test was applied to the simulation results generated using the Inferred Probabilities vs the Cell Variable models of isoform choice in Supplementary Figure 4 to test whether the distributions generated by different isoform choice models significantly differ.

| Data source | p-Value |
|---|---|
| H1 1 million reads | 0.99808 |
| H1 4 million reads | 0.981612 |
| H9 1 million reads | 0.989299 |
| H9 4 million reads | 0.997866 |

Table 9: Results of K-sample Anderson–Darling test, which tests whether multiple collections come from the same population. The test was applied to the simulation results generated using the Normal, Bernoulli and p=0.25 models of isoform choice to test whether the distributions generated by different isoform choice models significantly differ.
